# Raptor: A fast and space-efficient pre-filter for querying very large collections of nucleotide sequences

**DOI:** 10.1101/2020.10.08.330985

**Authors:** Enrico Seiler, Svenja Mehringer, Mitra Darvish, Etienne Turc, Knut Reinert

## Abstract

We present Raptor, a tool for approximately searching many queries in large collections of nucleotide sequences. In comparison with similar tools like Mantis and COBS, Raptor is 12-144 times faster and uses up to 30 times less memory. Raptor uses winnowing minimizers to define a set of representative *k*-mers, an extension of the Interleaved Bloom Filters (IBF) as a set membership data structure, and probabilistic thresholding for minimizers. Our approach allows compression and a partitioning of the IBF to enable the effective use of secondary memory.

## Background

The recent improvements of full genome sequencing technologies, commonly subsumed under the term NGS (Next Generation Sequencing), have tremendously increased the sequencing throughput. Within 10 years, it rose from 21 billion base pairs [1, 2] collected over months to about 400 billion base pairs per day (current throughput of Illumina’s HiSeq 4000). The costs for producing one million base pairs could also be reduced from many thousands of dollars to a few cents. As a result of this dramatic development, the number of new data submissions, generated by various biotechnological protocols (ChIP-Seq, RNA-Seq, etc.), to genomic databases has grown dramatically and is expected to continue to increase faster than the cost and capacity of storage devices can keep up. Ongoing projects like the 100,000 Genome Project [3] or the American 1,000,000 Genome Project [4] will easily produce data in the range of several petabases. This growth not only challenges the storage infrastructures and the processing pipelines of public databases because of the sheer data throughput, but also challenges algorithm engineers to improve the efficiency of sequence analysis pipelines and develop new strategies for compression, data parallelism, and concurrent computing.

The main task in analyzing NGS data is to search sequencing reads or short sequence patterns (e.g., read mapping and variant calling) or analyzing expression profiles in large collections of sequences (i.e. a database). Searching the *entirety* of such databases mentioned above is usually only possible by searching the metadata or a set of results initially obtained from the experiment. Searching (approximately) for specific genomic sequence in all the data has not been possible in reasonable computational time. The demand for solutions can be seen by the various attempts towards enabling sequence searches on large databases (see [5] for an overview). While the NIH SRA provides a sequence search functionality, the search is restricted to a limited number of experiments. Full-text indexing data structures, such as the FM-index, are currently unable to mine data of this scale. Word-based indices, such as those used by internet search engines, are not directly appropriate for edit-distance based biological sequence searches [6]. The sequence-specific solution CaBLAST [7] and its variants require an index of known genomes, genes or proteins, and thus cannot search for novel phenomena in raw sequencing files. In addition, none of these existing approaches are able to match a query sequence that spans multiple short sequences. This holds also true in the field of mapping-based metagenomic binning and quantitation where the relevant microbial databases grow about as fast as the sequence archives. The NCBI Refseq database of prokaryotic genomes contains about 30 GiB of sequence, still small enough to build an FM-index for the genomes, which takes about 24 h time and about 50 GiB memory [8]. However, including the draft genomes into the analysis increases the database to 380 GiB. Building a single search structure like an FM-index for this amount of data is infeasible.

### Related work

The problem of approximately searching queries in ultra-large databases has recently been addressed by several groups, focusing on different applications, but all using methods based on the *k*-mer content of the databases. In the field of alignment-free metagenomic analysis, which focuses on *k*-mer based analysis, the size of the data also becomes slowly prohibitive. For example, Kraken [9] needs ≈ 147 GiB RAM for indexing 380 GBases. For analyzing RNA-Seq data, some groups aimed at searching the raw files directly for a set of transcripts ([10] and shortly afterwards [11]). They propose novel solutions to the problem of searching a transcript of interest in all relevant RNA-Seq experiments. Up until recently, these searches were only based on the sequences itself; the tool REINDEER [12] is the first approach to also account for the sequence abundances.

As a benchmark, all three publications use a data set of 2, 652 RNA-seq sequencing runs of human blood, breast and brain tissue (a total of ≈ 6.5 TiB) in which they search for 214, 293 known transcripts. For a *single* query their methods need in the range of 2-20 minutes, which is a tremendous improvement and a speedup of orders of magnitude compared to previous methods. Although a breakthrough, the methods presented by the groups need 4 and 0.3-2 days for processing the above set of 214, 293 queries, respectively. Very recently, this time was improved by the Padro group with the tool Mantis in [13] to 82 minutes. Moreover, Bradley et al. [14] propose a Bloom filter based solution that can index about 170 TiB of (repetitive) raw sequence into an index of 1.5 TiB. However, searching, for example, 220 MiB of plasmid sequence takes 11 days using 8 cores. The same group followed up with a newer approach called COBS [6]. Finally, the construction time of an index build on top of 170 TiB of input data was further improved by the tool RAMBO [15] which only needs 14 hours on a cluster of 100 nodes. Taken together, all approaches are still very demanding in terms of memory consumption and run time.

### Our contribution

In this paper we propose a data structure, called *binning directory*, that can distribute approximate search queries based on an extension of the recently introduced Interleaved Bloom Filters (IBF) [8]. A *binning directory* combines a so called *x*-partitioned IBF (*x*-PIBF) with winnowing minimizers and probabilistic thresholding that takes into account the varying number of minimizers in each read. We present our implementation, called *Raptor*, discuss its capabilities and limitations, and compare it with the state-of-the-art methods Mantis and COBS. Our comparison shows that Raptor is up to 12-144 times faster than the competitors in answering approximate string searches with full sensitivity and a very good specificity. In addition, we only use a fraction of the memory and can further trade run time for main memory consumption.

## Results

Raptor stores a *representative* transformation of the *k*-mer content of the database that is divided up into a number of bins, typically a few hundred to a few thousand (see Methods section for details). The term *representative* indicates that the *k*-mer content could be transformed by a function which reduces its size and distribution (for example, using winnowing minimizers on the text and its reverse complement or using gapped *k*-mers). In this work we use (*w, k*)*-minimizers* for computing representative *k*-mers. A (*w, k*)-minimizer is essentially the lexicographically minimal *k*-mer of all *k*-mers and their reverse complements in a window of size *w*. The same transformation is applied to the *k*-mers of the query (see Methods and Figure 1 for an example and details). Raptor uses a set membership data structure, the *x*-PIBF, to retrieve *binning bitvectors* indicating whether a representative *k*-mer is in a bin or not. It then combines the binning bitvectors of all representative *k*-mers in a query into a *counting vector* and applies a thresholding step to determine the membership of a query in a bin.

**Figure 1.**
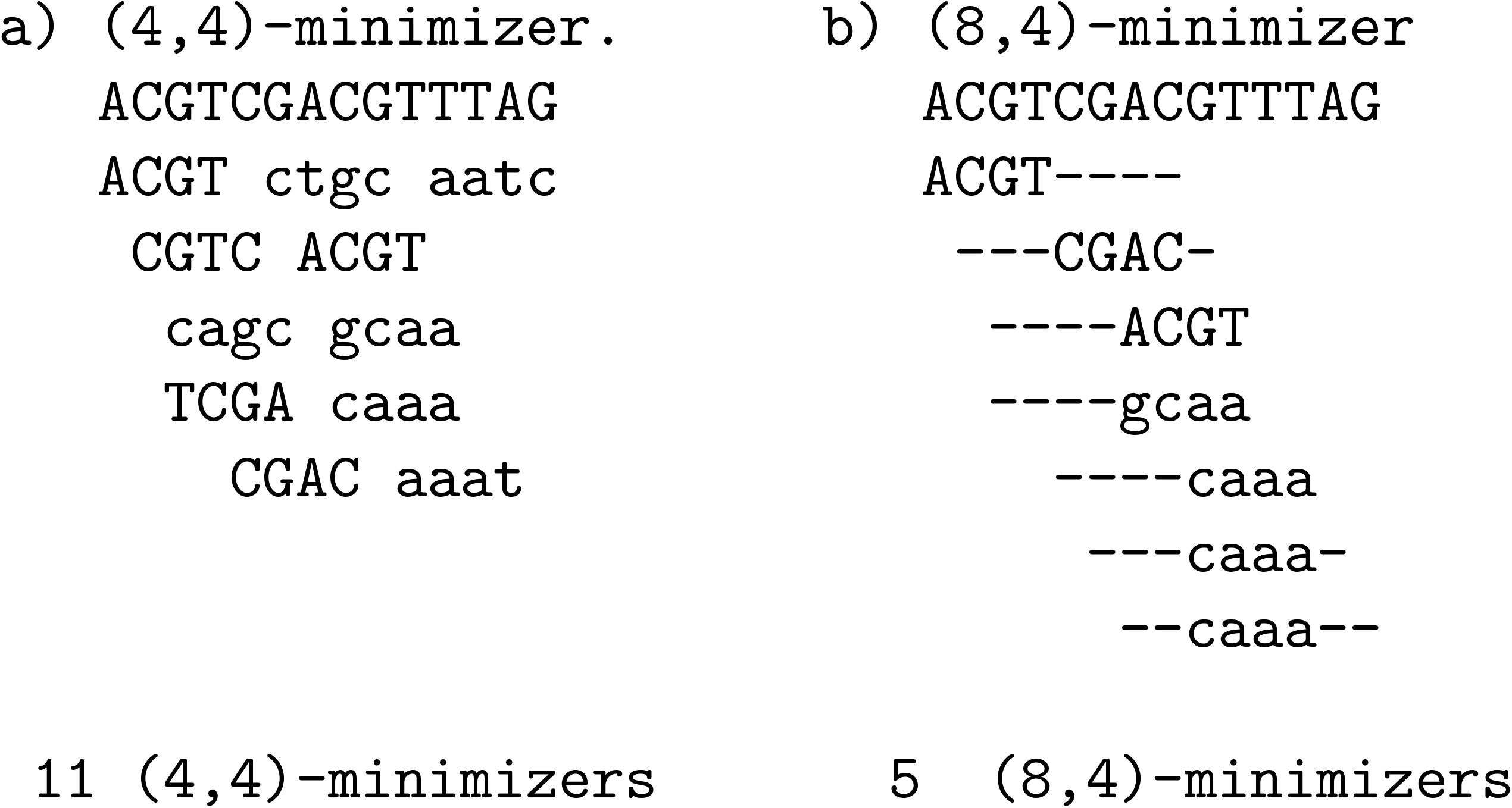
Examples of (*w, k*)-minimizers. a) and b) show the same *k* for different window sizes, the former with a window size *w* = *k* and the latter with a larger window of 8. Note that the reverse complemented sequences, shown in lower case, have to be read from right to left. The window width is indicated by a dash and the minimizing *k*-mer is placed within the window. Subsequent windows often share the same minimizer which we illustrated by showing those as well, although they are only stored once.

In the following, we report our computational experiments for Raptor. Firstly, we will use an artificial data set to discuss the limitations of binning directories, the impact of compression, the impact of the use of (*w, k*)-minimizers for different window sizes, and the time/space trade-off when using different partition sizes. We will also compare different binning directories with Mantis and COBS using this data set.

Secondly, we will evaluate Raptor using a real data set used by several groups to determine the membership of transcripts in RNA-Seq files [13] and compare Raptor with Mantis [13] and COBS [6].

All experiments were conducted on a Dell PowerEdge T640 with an Intel Xeon Gold 6248 CPU using 32 threads and 1 TiB of main memory. All file I/O was performed to and from a memory mapped file system (/dev/shm).

### Data sets

#### Artificial data set

We created a random DNA sequence of 4 GiB size and divided it into *b* bins which would correspond to *b* different genomes in, e.g., a metagenomic data set. Using the Mason genome variator [16], we then generated 16 similar genomes in each bin which differ about 1% from each other on average. This could be seen as bins containing the genomes for a very homologous species. The total sequence length is hence 64 GiB, however, containing *b* groups of highly similar sequences. Finally, we uniformly sampled a total of 2^20^ reads of length 100 bp from the genomes and introduced 2 errors in each read to simulate a sequencing experiment. On this data set we use (19, 19)-minimizers and (23, 19)-minimizers in conjunction with thresholds derived by the *k*-mer Lemma or our probabilistic thresholding for determining which bin contains the query. The value *k* = 19 was chosen to make random occurrences seldom (see Methods for a detailed discussion).

#### Real data set

In order to evaluate our method on real data, we took the data set from [13] which consists of 2, 568 RNA-Seq experiments. Similarly to [13], we excluded all experiments that have an average read length below 50*bp*, because reads shorter than that are rarely relevant in practice. Furthermore, this allowed us to test the minimizer approach with a broader window size. This left us with 1, 742 RNA-Seq experiments which have a size of around 6 TiB (gzipped FASTQ files). All tools were tested on this data set using *k* = 20, a value used in the competitors’ publications.

### Speed and space consumption of Raptor with (*w, k*)-minimizers

#### False positive (FP) count for different IBF sizes

A Bloom filter has a false positive rate depending on the ratio of stored elements to its size. For a fixed number of elements stored it holds that the less space we allocate, the more false positives will occur. In our experiment this will lead to overcounting *k*-mers and hence lead to false positive assignments of reads to bins. To evaluate this effect, we allocated IBFs of 1, 2, 4 and 8 GiB size and report the used RAM, construction time, search time and false positive bin assignments for *b* = 64 and *b* = 1, 024 bins for uncompressed and compressed vectors using *h* = 2 hash functions. Also, we use (19, 19)-minimizers and the traditional *k*-mer counting lemma threshold in one experiment, and (23, 19)-minimizers in conjunction with a new probabilistic threshold (see Methods) in a second experiment. The results are shown in Table 1.

**Table 1.**
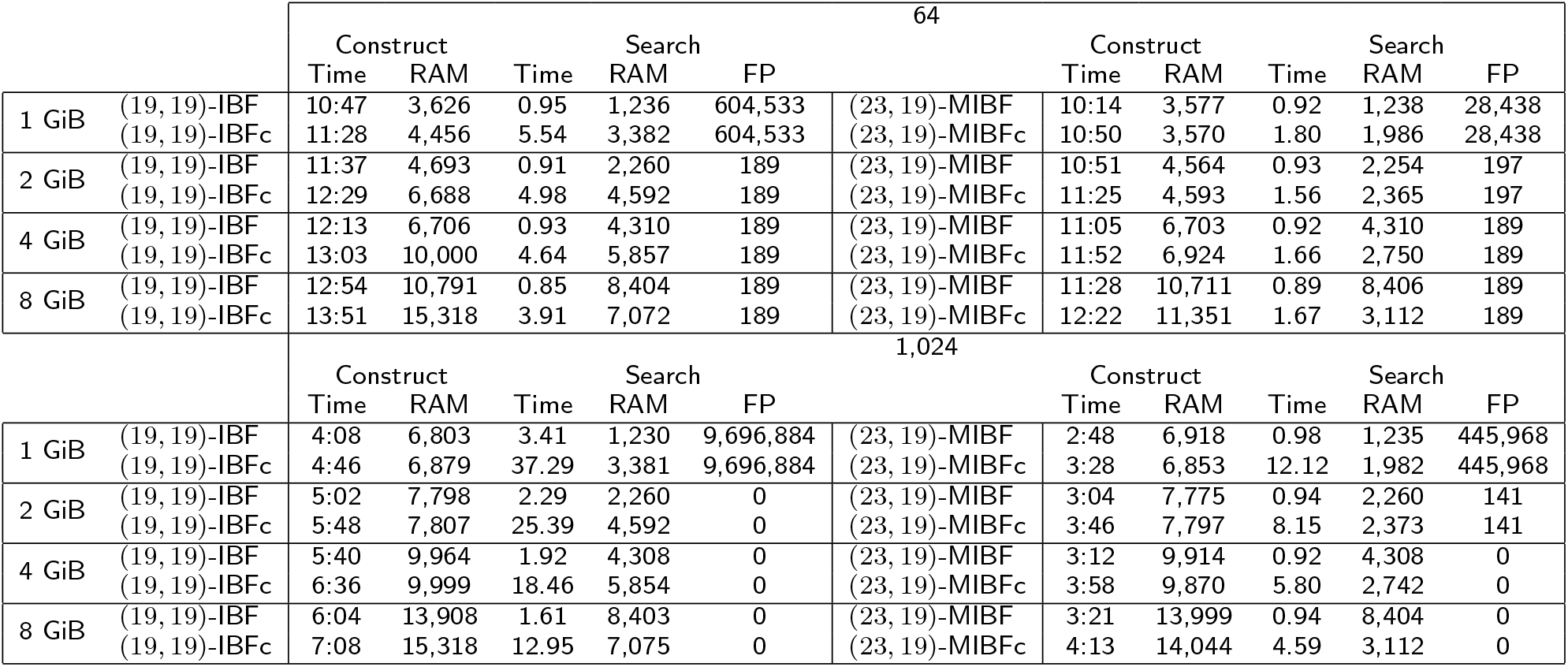
Run time and memory consumption of Raptor using differently sized IBF for *b* = 64 and *b* = 1, 024. On the left are the numbers for (19, 19)-minimizers (IBF), on the right for (23, 19)-minimizers (MIBF). Compressed versions are denoted by the suffix ‘c’. Construction times are in MM:SS, search times in SS.ss. RAM represents the memory peak in MiB during the construction and search, respectively. A total of 1,048,576 reads were processed, allowing for up to 2 errors. False positives (FP) are reads originating from bin *i* assigned to a bin *j ≠ i*, neglecting the fact that the read may match with bin *j* when allowing for 2 errors.

Our experiments show that allocating only 1 GiB for an IBF using (19, 19)-minimizers results in a high number of FP for all values *b* - we would have to conduct about 6 10^5^ and 9.6 10^6^ wrong verifications for the IBF for *b* = 64 and *b* = 1, 024, respectively. For (23, 19)-minimizers, the numbers are over one order of magnitude smaller (2.8 10^4^ and 4.4 10^5^). This is to be expected since we store a smaller set of representative *k*-mers. By doubling the size of the IBF, the number of false positives is already heavily reduced for (19, 19)-minimizers. Indeed, there are no more FP caused by the Bloom filter. Note that the 189 FP for *b* = 64 are reads whose minimizer composition actually occurs in a different bin than its original bin by chance. Since distributing the *k*-mers to more bins reduces the chance of the same minimizer composition being present in different bins, the FP count for *b* = 1, 024 is 0. For (23, 19)-minimizers, we can still see 197 − 189 = 8 FP induced by the Bloom filter for *b* = 64 and 141 for *b* = 1, 024. This indicates that the distribution of the (23, 19)-minimizers is not completely uniform or that our probabilistic threshold for the counting lemma introduced a few FP. In general, the effect of using minimizers on the FP rate is negligible. For larger sized IBF, no FP searches induced by the Bloom filter occur for both minimizer sets. The FP counts are obviously the same if we apply lossless compression to the bitvector.

#### Time and space usage for index construction

The construction time for *b* = 64 is between 11 and 13 minutes and for *b* = 1, 024 between 4 and 6 minutes. For 1, 024 bins, the wall clock time is smaller, since we can parallelize the construction (in chunks of 64) which is not possible for 64 bins. The space needed for construction is the size of the IBF and thread-local storage for the input sequences. In order to compress the IBF, both the uncompressed and compressed version must be in memory for a short amount of time, resulting in an increased memory peak.

For (23, 19)-minimizers, the construction time is generally lower since we insert fewer representative *k*-mers. While for *b* = 64, the times are comparable, the IBF can be built almost twice as fast for *b* = 1, 024 compared to (19, 19)-minimizers.

#### Time and space usage for the search

For *b* = 64, Raptor needs about 1 s to search for the 2^20^ reads for all IBF sizes. This holds true for both minimizer sets. Although Raptor searches fewer representative *k*-mers in case of the (23, 19)-minimizers, we need to compute the minimizers of the query beforehand, which is additional work. For *b* = 1, 024 we need between 2.2 s for a 2 GiB IBF and 1.6 s for a 8 GiB IBF. The increase for larger *b* is to be expected since we need to check for all bits in the binning bitvector. This takes longer for larger binning bitvectors. For (23, 19)-minimizers, we only see a slight increase and still need about 1 s. The benefit of querying fewer *k*-mers becomes pronounced and the IBF is up to twice as fast as the IBF for (19, 19)-minimizers.

When searching, it is also interesting how large the memory footprint is if we use compressed bitvectors. For *b* = 64 and (19, 19)-minimizers, we see for the IBF that compression actually *increases* the memory footprint until we use a IBF of 8 GiB. This means that the bitvectors are not sparse and that the space overhead of the compression algorithm outweighs the benefit of compressing the data.

In addition, the search time increases by a factor of about 4 for *b* = 64 and about 8-11 for *b* = 1, 024, which makes compression here unattractive.

This changes for (23, 19)-minimizers. For *b* = 64, the search time increases from about 1 s to only 1.6 s while we can compress the bitvector from 4.3 to 2.7 GiB or from 8.4 to 3.1 GiB. For *b* = 1, 024, the compression is similar since we store the same number of *k*-mers, but the run time increases by a factor of 5-8. This is due to the need to uncompress the 1, 024 bit long binning bitvector which takes longer than for the 64 bit long bitvector. Still, for (23, 19)-minimizers and smaller *b*, using compression offers an attractive time/space trade-off. For querying, we can observe that, in general, a sparser bitvector returns the results faster.

### Impact of the number of partitions on the speed

Finally, we investigated the impact of partitioning the IBF into *x* = 1, 2, 4, 8 parts. Since Raptor cannot di-rectly evaluate the counting vector for each read after having looked at one part, we need to store the intermediate results and check if we match the threshold after having counted the *k*-mers in all parts of the partition. To do this, Raptor allocates a buffer vector of size 10^7^ where each position holds a vector of *b* bits that is assigned to one of the reads. After counting the occurrences of the *k*-mers of a read in one partition, we can add the result to the vector and use the vector for the next batch of reads. We report on the construction and search time as well as on the maximum memory allocated by the resulting *x*-PIBF, for *b* = 64 and *b* = 1, 024 bins. We use an 8 GiB IBF in these experiments. The results are shown in Table 2.

**Table 2.**
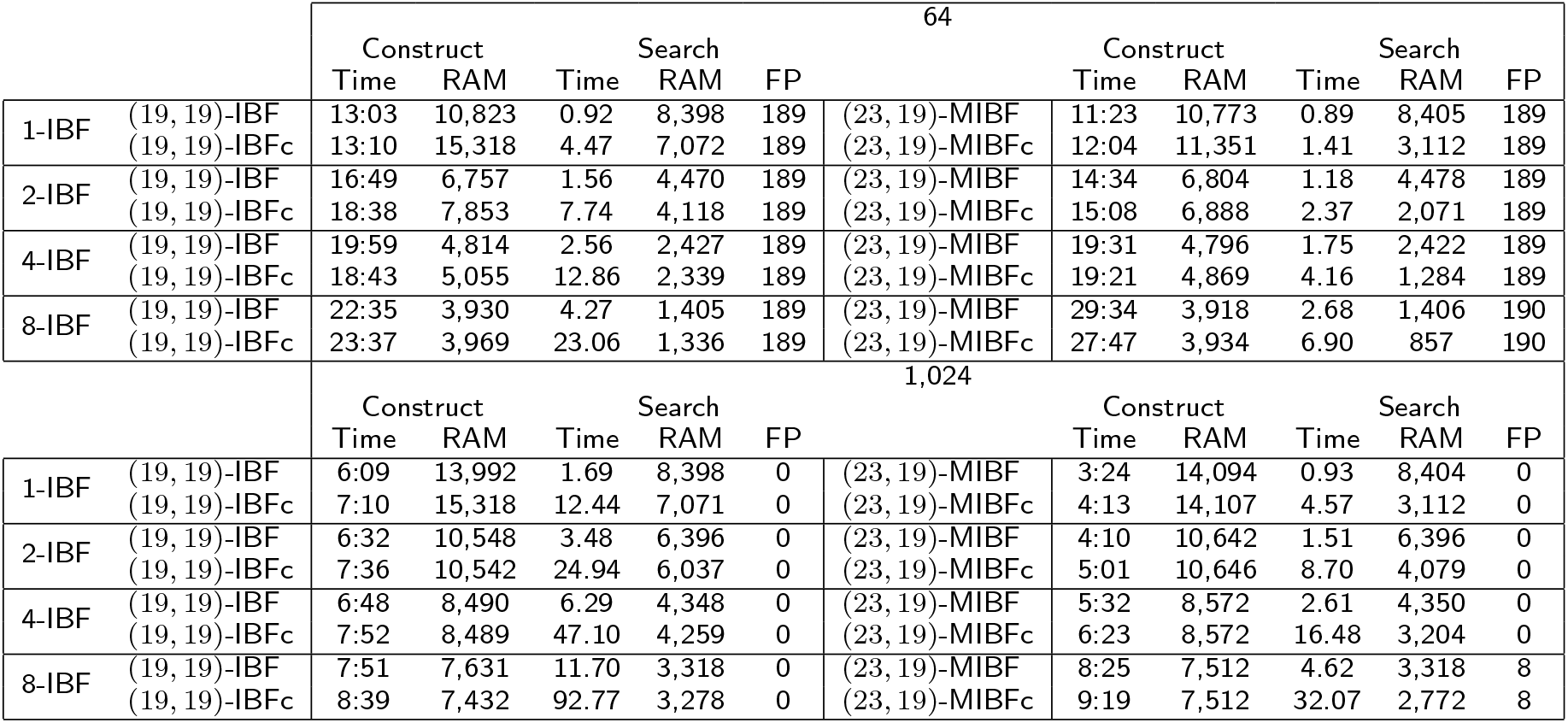
Construction and search time for partitioned IBF of size 8 GiB. The IBF is partitioned into 1 to 8 parts. On the left are the numbers for (19, 19)-minimizers (IBF), on the right for (23, 19)-minimizers (MIBF). Compressed versions are denoted by the suffix ‘c’. Construction times are in MM:SS, search times in SS.ss. RAM represents the memory peak in MiB during the construction and search, respectively. Raptor processes a total of 1,048,576 reads, allowing for up to 2 errors. False positives (FP) are reads originating from bin *i* assigned to a bin *j ≠ i*, neglecting the fact that the read may match with bin *j* when allowing for 2 errors.

In general, we observe for all minimizers that the construction and query times grow higher the more parts Raptor uses. For (19, 19)-minimizers, the build time increases from about 13 minutes to 23 minutes for *b* = 64, while for *b* = 1, 024 it increases from about 6 to 8 minutes. For *b* = 64, the search time for the IBF increases from about 1 s for a 1-PIBF to 4.3 s for a 8-PIBF. When using more parts, the run time increases, but the space needed to hold a single part in memory decreases. While we need 8 GiB memory to use a 1-PIBF, we only need 1.4 GiB if we use an 8-PIBF. When using (23, 19)-minimizers, we see similar trends for *b* = 64. Furthermore, like in the unpartitioned case, the search time is faster. Indeed, for an 8-PIBF we need only 2.68 seconds for the query and for an 8-PIBFc only 6.9 seconds while using only 857 MiB peak memory.

For *b* = 1, 024 and (19, 19)-minimizers, the search times for the IBF increases from 1.69 s to 11.7 s for a 8-PIBF. As before, compression is unattractive for this case, while it pays off for the (23, 19)-minimizer version.

In general, the construction time of Raptor’s index increases, the more parts we create, since we have to stream over our input *x* times and store *x* parts on the disk. However, we observe that this increase has a lower rate than the increase in the parts, as both constructing and querying a *x*-PIBF do take less than *x* times the time of a 1-PIBF. The reason for this is that we do not have to access the bitvector for *k*-mers that are not in the current part.

### Impact of probabilistic thresholding on false negatives

In this section we show that our probabilistic thresholding is crucial in avoiding false negatives. When we use (19, 19)-minimizers, the *k*-mer lemma ensures that we have no false negatives, but this is no longer true when using minimizers with *w > k*. This is apparent since the *number* and *distribution* of (*w, k*)-minimizers is sequence dependent and hence leads to a different threshold for each read. In the Methods section we describe how we derive a method to compute a threshold depending on the the parameters *w*, *k* and the number of minimizers a query has.

Indeed, Table 3 shows that a simple adaption of the *k*-mer Lemma for (23, 19)-minimizers would not work well. Instead of *k* we would use *w* as the length of the *k*-mer, i.e. 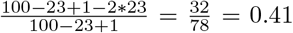. If we now require 41% of the (23, 19)-minimizers in a read to be present, it results in up to 2, 400 false negatives, which would not be acceptable. In contrast, our probabilistic threshold computation yields no false negatives for (23, 19)-minimizers. Tools like Mantis and COBS, which use a simple percentage cutoff, would suffer in a similar increase in false negatives if they used minimizers. However, they could use our results to adapt their thresholding. In our data set, the number of minimizers for each read ranges from 15 to 35 while our thresholds range from 4 to 13.

**Table 3.** False positives (FP) and false negatives for differently sized (23, 19)-MIBF using the adapted k-mer lemma. The resulting threshold is 41%. False positives are reads originating from bin *i* assigned to a bin *j i*, neglecting the fact that the read may match with bin *j* when allowing for 2 errors. False negatives are reads originating from bin *i* not assigned assigned to bin *i*.

|  | 64 |  | 1,024 |  |
| --- | --- | --- | --- | --- |
|  | FP | FN | FP | FN |
| 1 GiB | 309 | 796 | 1803 | 753 |
| 2 GiB | 189 | 1803 | 0 | 1696 |
| 4 GiB | 189 | 2270 | 0 | 2172 |
| 8 GiB | 189 | 2422 | 0 | 2308 |

For example, for a query of length 100 with 20 minimizers the threshold is 6, which is lower than 0.41*·*20 = 8.2. Using our threshold avoids falsely filtering out the query. In general, the percentage of minimizers that need to be present ranges from 26% to 38%. This shows that applying a single threshold is not sufficient.

### Comparison with other tools

In the following, we compare Raptor with the state-of-the-art tools Mantis [13] and COBS [6] using the artificial data set and a real data set of 1, 742 RNA-Seq experiments also used in [13] as described earlier. Note that the computational experiments for the real data set only use one thread for all tools, the same as it was done in the competitors’ publications. The effect of parallelization was tested using the artificial data set where we used 32 threads.

#### Artificial data set

We built an index over the artificial data set (separated in 64 and 1, 024 bins) with COBS and Mantis for a *k*-mer size of 19. Afterwards, we queried the same reads we have already searched with Raptor using BDs. Both COBS and Mantis consider a transcript found if the amount of *k*-mers found is more or equal to a given threshold. Instead of using the default threshold of 80 percent, we determined a threshold according to the standard *k*-mer counting lemma, which was 53 percent.

Moreover, we had to adapt our input for the index construction of Mantis by adding random quality scores to our FASTA files because Mantis only accepts FASTQ files as input. But even with this adaptation, Mantis, or more precisely the helper tool Squeakr, resulted in a segmentation fault for the artificial data set separated in 64 bins, thus we only present result for Mantis with 1, 024 bins.

As can be seen in Table 4, the construction of COBS and Mantis takes at least three times longer than for Raptor. Furthermore, searching with Raptor only needs a fraction of the space (about 5-8 GiB vs. 20 GiB) COBS and Mantis need, while being as accurate. The most striking difference is the search time. For (19, 19)-minimizers, Raptor needs between 0.9 s for *b* = 64 and 1.6 s for *b* = 1, 024. This is about 144 times faster than COBS and (for *b* = 1, 024) about 30 times faster than Mantis.

**Table 4.**
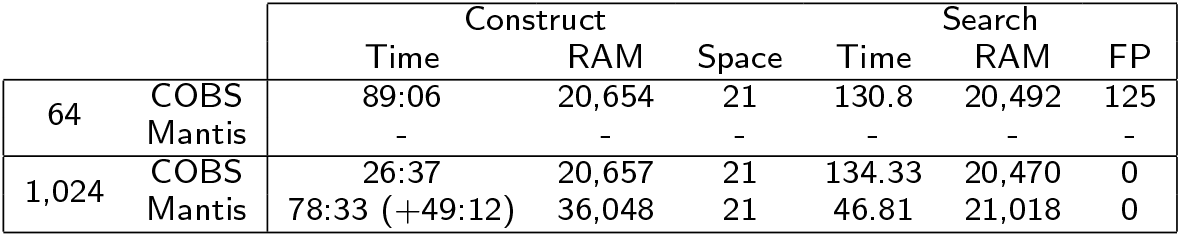
Comparison COBS and Mantis for the artificial data set with 64 and 1,024 bins. Construction times are in MM:SS, search times in SS.ss. The construction time in brackets for Mantis is the additional time the preprocessing tool Squeakr needs. The used disk space is in GiB, the maximum RAM in MiB. All approaches are built for *k* = 19.

|  |  | Construct |  |  | Search |  |  |
| --- | --- | --- | --- | --- | --- | --- | --- |
|  |  | Time | RAM | Space | Time | RAM | FP |
| 64 | COBS | 89:06 | 20,654 | 21 | 130.8 | 20,492 | 125 |
|  | Mantis | - | - | - | - | - | - |
| 1,024 | COBS | 26:37 | 20,657 | 21 | 134.33 | 20,470 | 0 |
|  | Mantis | 78:33 (+49:12) | 36,048 | 21 | 46.81 | 21,018 | 0 |

#### Real data set

We built an index over the real data set with COBS, Mantis, and Raptor using a binning directory for a *k*-mer size of 20 (since this value was used in [13], see Additional File 2). For Raptor, we created two versions, one using a binning directory with an IBF with (20, 20)-minimizers and one version using a binning directory with an IBF with (40, 20)-minimizers. Mantis uses a cutoff in order to sort out low-frequency *k*-mers that probably resulted from sequencing errors [13]. In order to be comparable, Raptor applied the same cutoffs for both versions of the binning directories. The results are shown in Table 5.

**Table 5.** Comparison of Raptor, COBS and Mantis. Construction times are in minutes. The used disk space and the maximum RAM are given in GiB. All approaches are built for *k* = 20.

|  | Construct |  |  |
| --- | --- | --- | --- |
|  | Time | RAM | Space |
| (20, 20)-IBF | 34 | 8.1 | 8 |
| (40, 20)-IBF | 2 | 0.9 | 0.8 |
| COBS | 4,620 | 702.6 | 4,265 |
| Mantis | 135 | 29.7 | 17 |

Raptor’s construction time of the binning directory is faster than Mantis and COBS and, when using (40, 20)-minimizers, the space consumption drops to only five percent of that of Mantis. COBS’s construction time and space consumption is nowhere near the other two applications, because COBS has no preprocessing step and does not use cutoffs to filter out erroneous *k*-mers. Therefore, further comparisons to COBS are omitted.

In order to compare the query times, three differently sized sets (100, 1,000, and 10,000 transcripts) were used. Each set was created by randomly picking human gene transcripts. The lengths varied between 46 bp and 101, 518 bp.

The false positive rate (FPR) was determined by comparing Raptor’s results to Mantis, assuming Mantis correctly finds all experiments as it claims to be exact. A similar evaluation was applied in [13]. Also, the definition of a found transcript is based on the evaluation of [13]. Therefore, both Mantis and Raptor consider a transcript found in an experiment if 80% of its representative *k*-mers are found.

As shown in Table 6, Raptor using a BD with (40, 20)-minimizers is significantly faster (12-58 times) than Mantis and uses only a fraction of main memory while still being specific with a low FP rate of about 0.017. Even when using (20, 20)-minimizers, Raptor outperforms Mantis in space and time consumption.

**Table 6.** Comparison of Raptor and Mantis. Search times are in seconds, RAM is given in GiB. All approaches are built for *k* = 20.

|  |  | Search |  |  |
| --- | --- | --- | --- | --- |
|  |  | Time | RAM | FPR |
| 100 transcripts | (20, 20)-IBF | 7 | 8 | 0.025 |
|  | (40, 20)-IBF | 1 | 0.8 | 0.015 |
|  | Mantis | 12 | 18.7 | 0.0 |
| 1,000 transcripts | (20, 20)-IBF | 10 | 8 | 0.03 |
|  | (40, 20)-IBF | 1 | 0.8 | 0.016 |
|  | Mantis | 30 | 19 | 0.0 |
| 10,000 transcripts | (20, 20)-IBF | 46 | 8 | 0.031 |
|  | (40, 20)-IBF | 4 | 0.8 | 0.017 |
|  | Mantis | 232 | 23.3 | 0.0 |

## Discussion

In this paper we presented an approach to answer approximate string queries using a *representative* set of *k*-mers of the database and query. We stored a set of (*w, k*)-minimizers as representative *k*-mers of the database in a binning directory using a partitioned, interleaved Bloom filter.

Binning directories could be used in various settings which we discuss below.

### Using BDs for metagenomic profiling

Tools like Kraken [9] or Centrifuge [17] perform metagenomic binning by querying the *k*-mer content of genomes using NGS reads and inferring the presence or absence of organisms in the sample using the taxonomy of a phylogenetic tree.

Hence, we could use BDs for a classification based on taxonomic levels (e.g., species, genus,…) or assembly level, and group the reference genome sequences accordingly. Using the counts for *k*-mers given by the BD, we can infer the composition of a metagenomic sample. Indeed, [18] already applied this idea as described in [8] for this task.

### Using BDs for querying file content

Another application for BDs which we also used in one benchmark is to query all existing human RNA-Seq files in the SRA for the presence of transcripts. For this application, the bins would be defined by the respective file content. We would expect that the effective text size *n*(*k*) is considerably less than 5 TBases since we sample from human genes. Of course, this application scenario is not limited to RNA-Seq files.

### Using BDs for read mapping

In the context of read mapping, we can use the BD as follows. The database would be the reference genome(s) we want to map our reads against. Assume we have partitioned them into bins such that similar parts of the genomes are placed within the same bin. In the context of metagenomics analysis, this could be achieved by using a taxonomic tree (see also [8]); alternatively, the sequences could be clustered based on similarity. The sequences in the bins could then be indexed using a compressed suffix array or other suitable indices and the BD can be used to distribute the approximate searches.

### Possible extensions

Currently, Raptor stores all representative *k*-mers, even if some representative *k*-mers in the reference data set are ubiquitous, i.e. they appear in all or almost all, e.g., 95%, bins. While some approaches, like Mantis, exclude certain *k*-mers from consideration, one could instead exclude them from the IBF and store them in a small lookup table. Whenever such a *k*-mer is queried, we can increase the counters on all bins and save the lookups in the IBF. This might reduce the size of the IBF and speed up the search time.

While not shown in this paper, the update operation on an IBF was already used in DREAM-Yara [8] and ganon [18]. Adding data is trivial since we just need to set the corresponding bits in the *x*-PIBF. For removing data from the *x*-PIBF we need to clear and rebuild the *affected* bins of the update.

## Conclusion

In conclusion, we presented a novel, versatile, fast, and memory efficient data structure for *k*-mer based analysis of large sets of partitioned sequences using binning directories. Our implementation, Raptor, is ready for secondary memory use and its data structures can be efficiently compressed if the used bitvector is sparse. Furthermore, we showed that the concept of (*w, k*)-minimizers allows to effectively reduce the set of representative *k*-mers without sacrificing specificity nor sensitivity by applying our probabilistic thresholding. Raptor outperformed the state-of-the-art tools Mantis and COBS in both run time and space consumption. The use of (40, 20)-minimizers was able to reduce the memory footprint of our method from 8 to 0.9 GiB for the RNA-Seq data set introduced in [13], which is about one order of magnitude less compared to Mantis (*≈* 19-23 GiB). Using (*w, k*)-minimizers, the run time was better by factors between 12 and 144 compared to Mantis and COBS which enables completely new ways for analyzing large sequencing archives in ways that were not possible before. Raptor and binning directories are available in the SeqAn library [19] of efficient data types and algorithms.

## Methods

### Binning directories

In the following, we give a more formal definition of binning directories and explain how we solve current bottlenecks.

#### (w, k)-minimizers

In this work we introduce the concept of (*w, k*)*-minimizers* for computing representative *k*-mers. In Figure 1, we show an example for this concept. The reverse complement sequence is denoted in lower case, and we used the lexicographically smallest *k*-mer for clarity. In practice, this leads to a skewed distribution of minimizers which can be corrected by, for example, applying an XOR operation with a random value to each *k*-mer hash value before taking the numeric minimum (see [20] for a discussion). In the Figure you see a short example of a) ungapped 4-mers in a window of size 4, which means we take the lexicographically smallest of the *k*-mer and its reverse complement as the minimizing *k*-mer. The second case b) shows the minimizing 4-mers for a window of size 8. We form the minimum of all *k*-mers and their reverse complement in this window. We denote the window span with ‘-’ and place the minimizing *k*-mer at the respective window position. For the properties and the size of the data we will handle, *k* will usually be in the range of 16-32.

#### Effective text size and ratio

In general, Raptor assumes that we have a collection of str s {*T*_*j*_} over an alphabet Σ, with a total length *n* = Σ_*j*_ *|T_j_|*. Raptor stores the *k*-mer content of {*T*_*j*_} or a *representative* transformation of it. Raptor uses (*w, k*)*-minimizers* for computing the set of representative *k*-mers.

To capture the repetitiveness of the {*T*_*j*_}, we define the *effective* text length *n*(*k*) as the number of distinct, *representative k*-mers in all the *T*_*j*_. In order to store the set of texts {*T*_*j*_}, we further assume that we have divided the *T*_*j*_ into *b* bins *B*_*i*_ (mind that a single *T*_*j*_ itself could be partitioned into several bins without many adaptations). For the strings in a bin *B*_*i*_, we denote the set of representative *k*-mers with *B_i_*(*k*) and the effective text length with *n_i_*(*k*) as the number of representative *k*-mers of the strings in *B*_*i*_, i.e. the cardinality of *B*_*i*_(*k*).

Dividing the strings into bins could result in a large or small intersection of their representative *k*-mer content, depending on the method. To capture this, we define the *effective text ratio r*_*i*_(*k*) as 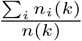. The effective text ratio is a measure of how well we have partitioned our *k*-mer content into the bins. Ideally it is 1 and in the worst case it is *b*. We want to point out that the effective text length *n*(*k*) is a crucial measure for the problem of indexing large genomic text collections. For example, [14] compute an index for 170 TiB of sequence data. However, this data set is quite repetitive since its effective text length *n*(31) is only 6·10^10^.

For our artificial data set we give the effective text ratio *r*(*k*) for different *k* for both 64 and 1, 024 bins in Table 7.

**Table 7.** Effective text ratios *r* (*k*) for 64 and 1, 024 bins and different values of *k* for the artificial data set. Values are rounded.

| b,k | 12 | 13 | 14 | 15 | 16 | 17 | 18 | 19 | 20 |
| --- | --- | --- | --- | --- | --- | --- | --- | --- | --- |
| 64 | 62.84 | 40.52 | 14.19 | 3.96 | 1.57 | 1.13 | 1.03 | 1.01 | 1.00 |
| 1,024 | 225.98 | 62.19 | 15.92 | 4.08 | 1.58 | 1.13 | 1.03 | 1.01 | 1.00 |

One can see that we need a *k*-mer size of at least 16 to achieve an effective text ratio under 2. For *k*≥ 19 the effective text ratio is near 1 which means that most *k*-mers in the bins are unique. For this reason we used *k* = 19 in our experiments.

#### Binning directory

We define a *binning directory (BD)* for the text collection {*T*_*j*_} divided into *b* bins *B*_*i*_ as a data structure that returns the counts of the representative *k*-mers in the query multiset *I*(*k*) for each bin *B*_*i*_. In this work a binning directory uses a set membership data structure, namely the *x*-PIBF, that returns the bin membership as a (compressed) bitvector which we call the binning bitvector. The BD then combines the binning bitvectors into count vectors. Our Tool Raptor uses (probabilistic) thresholding to determine whether a query is in a bin or not.

Implementing a simple version is indeed not difficult. The problems lies in the fact that the effective text size *n*(*k*) can be very large, i.e. 10^10^ to 10^12^. For example, the metagenomics data set used in [8] contains about 2·10^10^ different 19-mers. A naive implementation that stores all those 19-mers in a hash table containing the binning vectors for 1, 024 bins would need about 40 TiB (an open addressing hash table with about 4·10^10^ entries, each pointing to a 1, 024 bit bitvector). Hence, the challenge is implementing the BD in a more space efficient way while maintaining a fast run time.

We approach this problem in two ways in this paper. For implementing a binning directory, we adapted the IBF data structure presented in [8] to work well on secondary memory. We call it the *x*-partitioned IBF (*x*-PIBF). Secondly, we employ (*w, k*)-minimizers to reduce the number of representative *k*-mers significantly while still accurately answering the question in which bins a query can occur.

#### Answering a query with Raptor

Answering a query includes the retrieval of the binning bitvectors and the counting of *k*-mers to determine the bins to be searched. Using a *x*-PIBF, Raptor has to compute *h* hash functions, retrieve *h* sub-bitvectors and compute a bitwise AND. We can use a standard bitvector of size *n* that uses *n* bits, or the compressed bitvector of the SDSL [21] that uses approximately 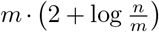 bits, where *m* is the number of bits set and *n* the length of the bitvector.

For counting, Raptor has to traverse the binning bitvector of size *b* and increment counters for each bit set to 1. To speed up this crucial step we used, for uncompressed bitvectors, the lzcount intrinsic operation which counts the number of leading zeros in a 64 bit word. This accelerated the bin counting step by a factor of almost 2 compared to the individual checking of each bit.

A further speed up is possible once the AVX512 SIMD extensions are available on standard computers (already possible for Intel’s Skylake processor). These optimizations cannot be directly applied to compressed bitvectors.

Having the counts, we apply a thresholding according to the original *k*-mer counting lemma [22] or according to a probabilistic model for (*w, k*)-minimizers.

##### Lemma 1

*For a given k and number of errors e, there are k*_*p*_ = ∣*p*∣ −*k* + 1 *many k-mers in p and an approximate occurrence of p in T has to share at least t* = (*k*_*p*_ − *k · e*) *k-mers.*

Hence, if the count exceeds the threshold for the bin, we report the pattern to occur in this bin, otherwise not. This approach is depicted in Figure 2. However, using minimizers makes the direct application of Lemma 1 impossible for *w > k*. We present a solution in the next section.

**Figure 2.**
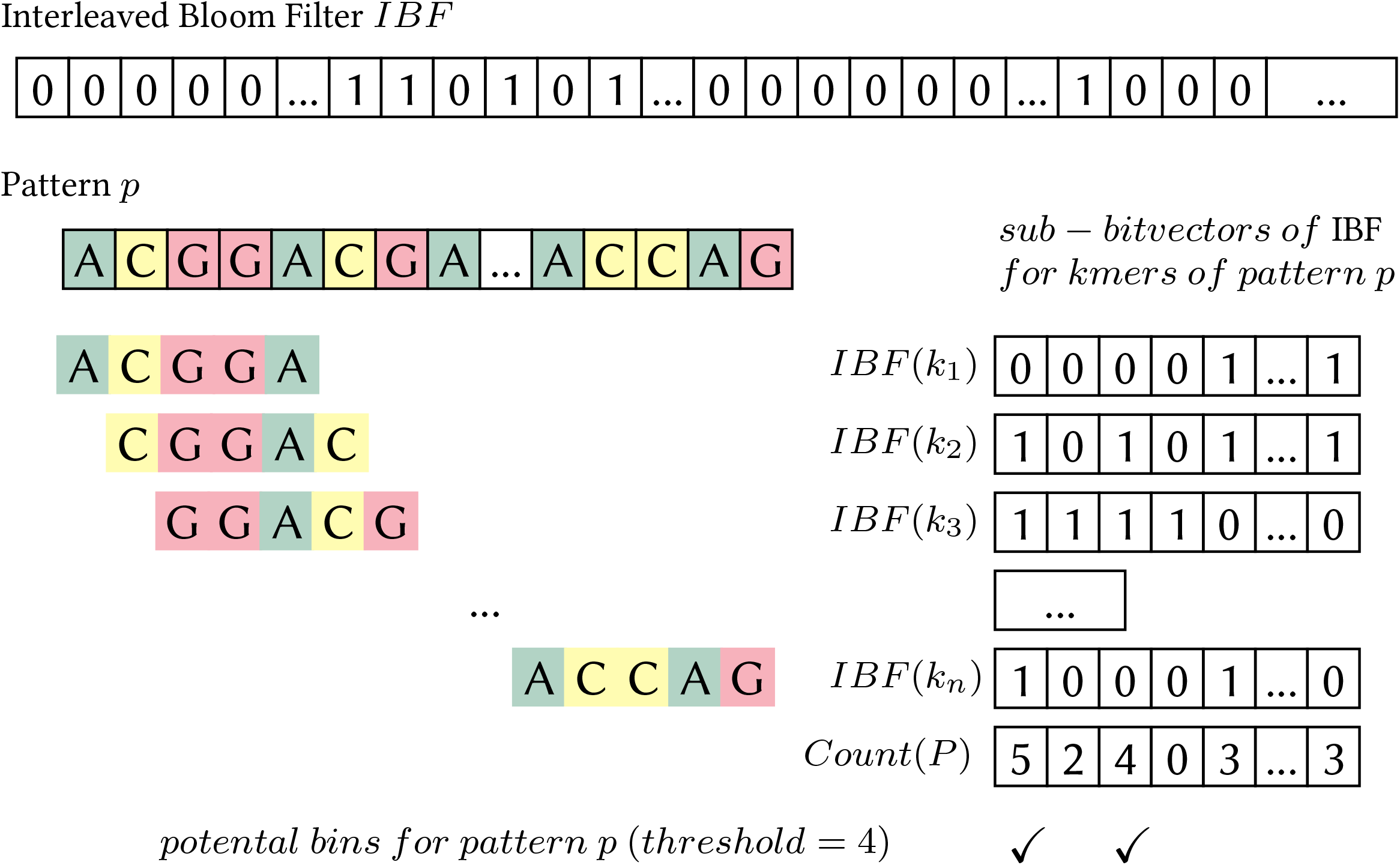
Binning directory in conjunction with the *k*-mer Lemma. Bins with a counter greater than or equal to the threshold (in this case 4) need to be validated for *p*.

### Probabilistic thresholding

Lemma 1 works only for (*k, k*)-minimizers. It does not hold for general (*w, k*)-minimizers. The latter is apparent since the *number* and *distribution* of (*w, k*)-minimizers is sequence dependent and hence leads to different thresholds for each read.

This problem is exemplified in Figure 3. The examples show that an error does not only *directly* invalidate the minimizers covering the error position but also *indirectly* affects minimizers *not* covering the error position, resulting in a different count of minimizers.

**Figure 3.**
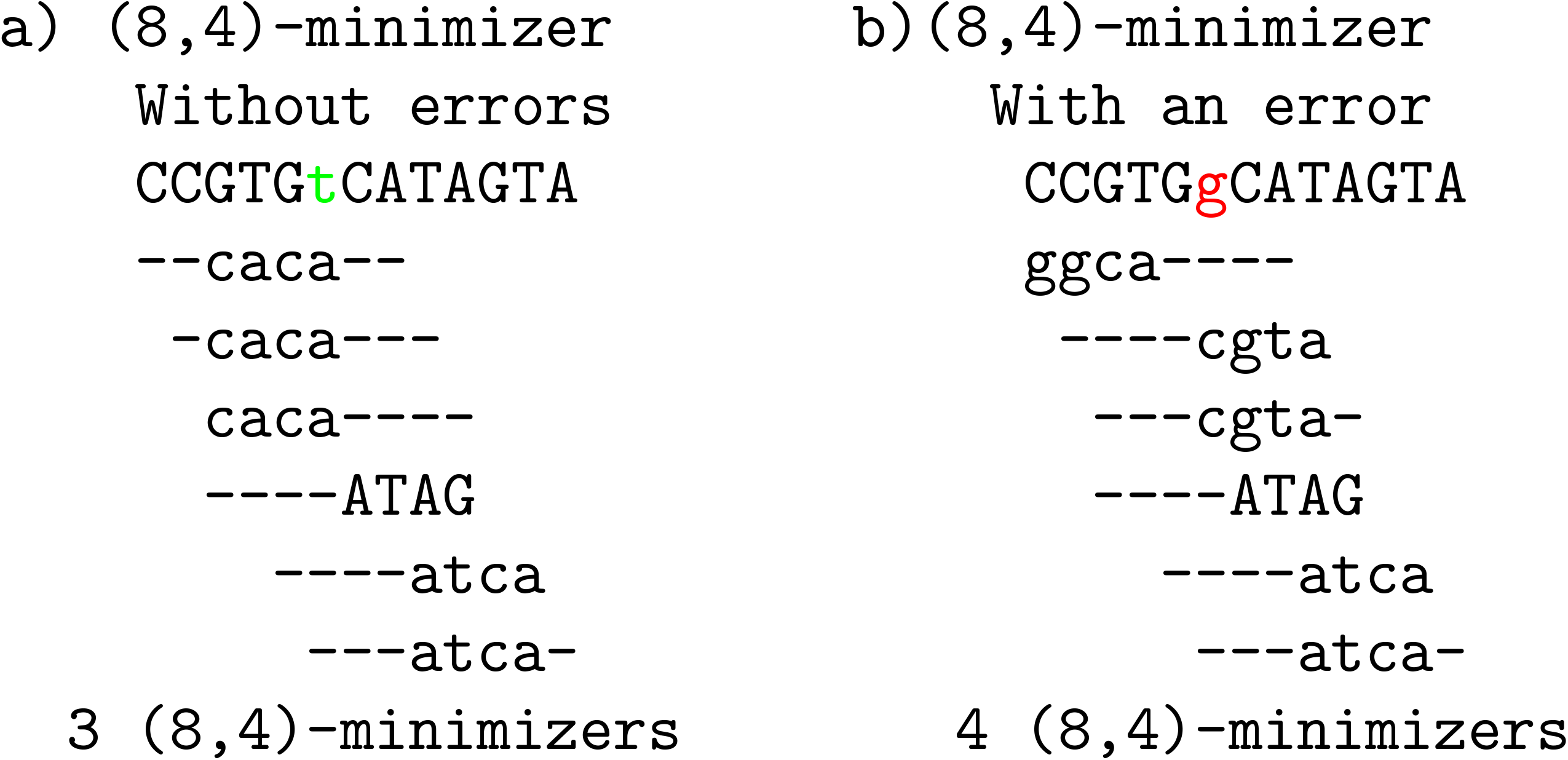
Impact of a direct error on the number of *k*-mers. The sequences a) and b) represent the same sequence without and with an error at position 6 replacing a T with a G, respectively. The sequence in a) has 3 minimizers, one of which (*caca*) is destroyed by the error position. Hence, we could assume that a sequence with one error at this position has a count of 2. However, introducing the error by replacing T with G has the effect that the first window now has a different minimizer not covering the errorposition (*ggca*) and hence b) still has a minimizer count of 3. Thus, a) and b) would be wrongly deemed not matching with 1 error.

Taking these indirectly destroyed minimizers into consideration, there are several ad hoc ways to compute the threshold. The first is to adapt Lemma 1 such that we compute the threshold as follows: For a given *k*, *w* and number of errors *e*, there are *w*_*p*_ = ∣*p*∣ −*w*+1 many windows in *p* and if we take the multiplicity of the minimizers into account, an approximate occurrence of *p* in *T* has to share at least *t* = (*w*_*p*_ − w · *e*) minimizers, i.e. we replace *k* with *w*. However, this leads to low thresholds. The threshold in Figure 3 *a*) would be negative, i.e. 13 − 8 + 1 − 8 = −2, and thus useless for filtering.

Another way to compute an individual threshold is to repeat the following two steps for each error: 1) compute the minimizer coverage of a query *p* (counting each minimizer only once) 2) One maximum coverage position is chosen and the minimizers covering this position are removed. The overall number of removed minimizers is subtracted from the number of minimizers to obtain the threshold *t*. This works better than the first approach, but is quite time-consuming to compute.

We can show that a simple probabilistic model yields thresholds that are on average much better than the above ad hoc methods and removes the need to compute an individual threshold for each read. Instead, the threshold can be looked up in a table given the number of minimizers of a query and the parameters (*w, k*) and hence the filtering speeds up considerably. The method models the occurrences of (*w, k*)-minimizers within the windows and how they affect each other (see Figure 3 for an example). Our method models the distributions of the minimizer occurrences and how they affect each other. Given a threshold *τ* for the cumulative probability of *d* minimizers being affected by errors, we compute a value for the number of minimizers not affected by errors. Details can be found in Additional File 1.

### *x*-Partitioned Interleaved Bloom filter

Finally, we propose our last contribution, the use of an *x*-partitioned interleaved Bloom filter (*x*-PIBF) in binning directories, which are an extension of the IBF proposed in [8].

An IBF for *b* bins combines *b* standard Bloom filters [23]. A Bloom filter is simply a bitvector of size *n* and a set of *h* hash functions that map a value, in our case a representative *k*-mer, to one of the bit positions. A value is present in the Bloom filter if all *h* positions return a 1. Note that a Bloom filter can give a false positive answer. However, if the Bloom filter size is large enough, the probability of a false positive answer is low. A Bloom filter of size *n* bits with *h* different hash functions and *m* elements inserted has a probability of giving a false positive answer of approximately

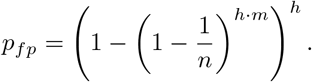

For this reason, we have to allocate sufficient space such that *p*_*fp*_ does not become too large. Still, the problem of using a simple Bloom filter is that it does not point us to the binning bitvectors. To alleviate the problem, the IBF uses *b* Bloom filters (one for each bin) with identical hash functions and then interleaves their bitvectors. Putting it differently, this means that it replaces each bit in the Bloom filter by a (sub)-bitvector of size *b*, where the *i*-th bit “belongs” to the Bloom filter for bin *B_i_*. The resulting IBF has a size of *b · n*. When inserting a *k*-mer from bin *B*_*i*_ into the IBF, it computes all *h* hash functions which point to the position of the block where the sub-bitvectors are and then sets the *i*-th bit from the respective beginnings. Hence, the IBF effectively interleaves *b* Bloom filters in a way that allows us to easily retrieve the binning bitvectors for the *h* hash functions. When querying in which bins a *k*-mer can be found, we retrieve the *h* sub-bitvectors and apply a logical AND to them which results in the required binning bitvector indicating the membership of the *k*-mer in the bins. The procedure is depicted in Figure 4.

**Figure 4.**
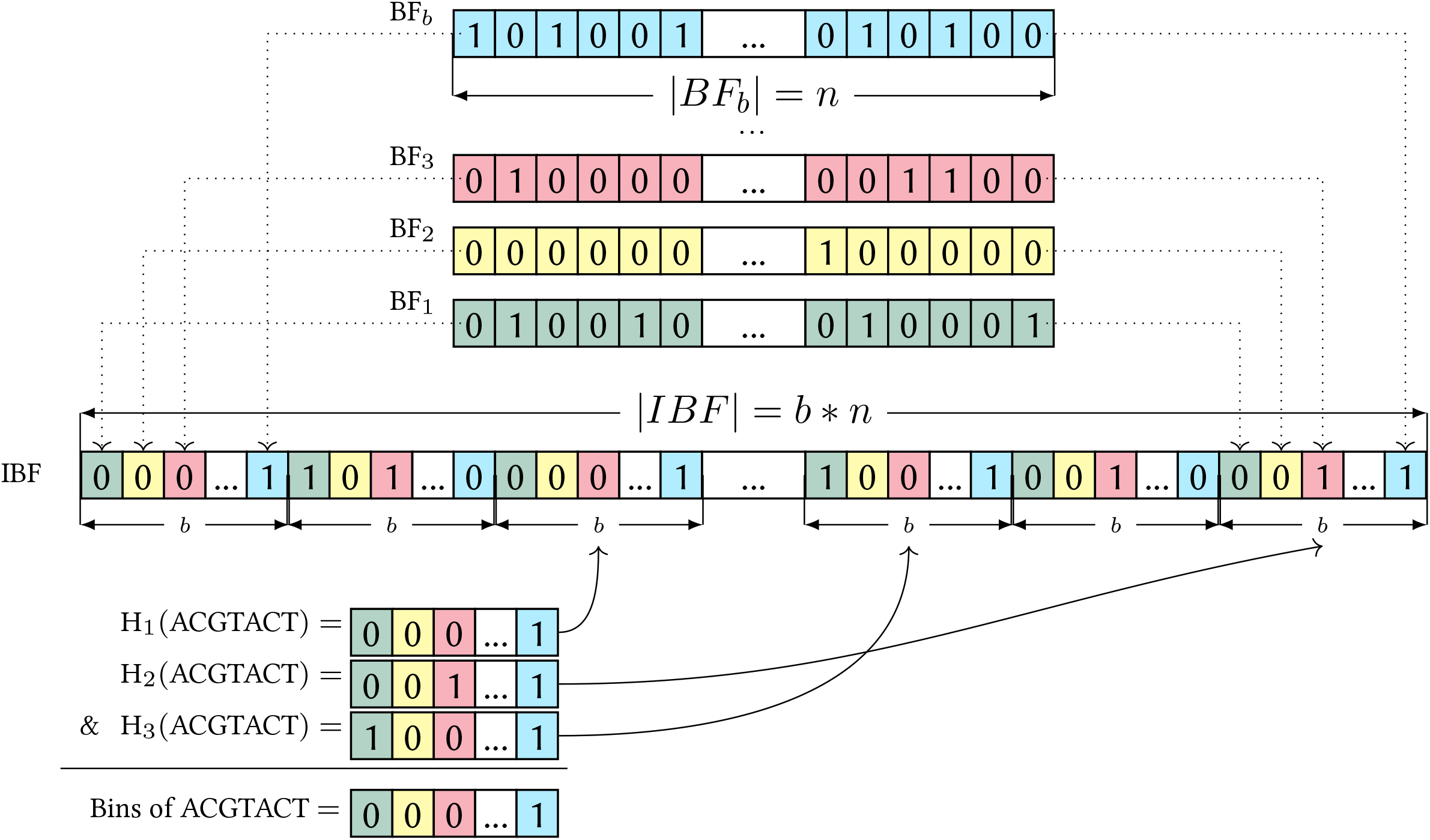
Example of an IBF. Differently colored Bloom filters of length *n* for the *b* bins are shown in the top. The individual Bloom filters are interleaved to make an IBF of size *b×n*. In the example we retrieve 3 positions for a *k*-mer (ACGTACT) using 3 different hash functions. The corresponding sub-bitvectors are combined with a bitwise & resulting in the needed binning bitvector.

Finally, the binning bitvectors are summed up to obtain the count vectors. For this, we allocate *t* many counters, each with *b* entries, where *t* is the number of threads used. These counters are reused as we process the reads in parallel.

#### Partitioning the IBF

If the set of stored *k*-mers is very large or if we want to achieve a very low false positive rate of the IBF, it might be too big to keep in the main memory. For those cases we implemented the *x*-partitioned IBF (*x*-PIBF) where we partition the set of stored *k*-mers into *x* parts as follows:

For a *x*-PIBF of size *s* bits, we create *x* parts, each with *[s/x]* bits. Then we partition the *k*-mers based on the first *q* characters with *q* ≥ [log_*σ*_ (*x*)], where *σ* is the alphabet size. We count the *q*-mer frequencies of all *k*-mers and assign them as evenly as possible to the *x* parts (see Table 8 for an example). The counting step can be omitted, in which case we assume a uniform prefix distribution.

**Table 8.**
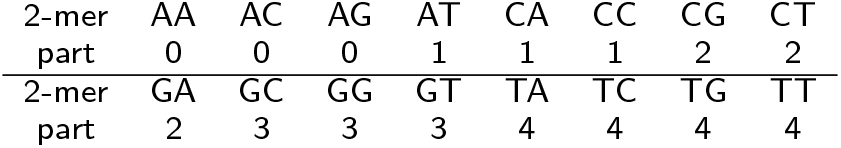
Example of the assignment of *q*-mers to x=5 partitions. Given a DNA alphabet (*σ*=4) and x=5, we have to distribute the 16 possible *q*-mers evenly to the 5 parts. In this example we assume a uniform distribution of the *q*-mers.

Finally, we have to adapt our hash functions such that all *h* hash values for a *k*-mer lie in the same part of the *x*-PIBF. This can easily be done by storing offsets in a *q*^*σ*^ large table and adding those to the hash values for an IBF of size *[s/x]*.

If we query a set of *k*-mers, we load the first part of the *x*-PIBF into memory, stream over all *k*-mers counting the relevant ones for this part and ignoring the others. Then we repeat this for all other parts after loading them. In the Results section we report on the time/memory trade-off.

### Compressing bitvectors

Binning directories use a large bitvector containing all the binning bitvectors for all representative *k*-mers. In this work we also allow the use of a compressed bitvector implementation from the SDSL [21]. While a standard bitvector of size *n* uses *n* bits, the compressed bitvector of the SDSL uses approximately 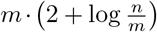 bits, where *m* is the number of bits set, and *n* the length of the bitvector. Note that while this reduces the space consumption, it increases the access time which we will discuss in the experiments.

To construct a compressed bitvector, we first have to create the *entire* uncompressed bitvector and then compress it. This means that both the uncompressed and compressed bitvectors have to be in main memory at some point during construction which increases the memory footprint during construction while reducing the memory requirements when using the bitvector. A main property of the compressed bitvector is that it is immutable. If we want to change a bit after the vector is constructed, we need to change the bit in the uncompressed bitvector and reconstruct the compressed bitvector. Since decompression for the compressed bitvector is not supported by the SDSL, we also need to store the uncompressed bitvector on disk to enable future updates of the IBF. Nevertheless, we need to have the whole bitvector initially in memory which might pose a problem. This problem can be solved elegantly using the partitioning of the IBF as proposed before.

## Acknowledgements

The authors acknowledge the support of the de.NBI network for bioinformatics infrastructure, the Intel SeqAn IPCC, the Max-Planck society and the IMPRS for Biology and Computation (BAC).

## Abbreviations

BD: Binning Directory
FP: False Positive
FPR: False Positive Rate
IBF: Interleaved Bloom Filter
MIBF: Minimizer Interleaved Bloom Filter
PIBF: Partitioned Interleaved Bloom Filter
SDSL: Succinct Data Structure Library

## Availability of data and materials

- Project name: Raptor
- Project home page: https://github.com/seqan/raptor
- Archived version: 1.0.0
- Operating systems: Unix, OSX
- Programming language: C++
- License: BSD-3-Clause

## Authors’ contributions

ES designed and wrote the software and conducted the experiments on the artificial data set. SM designed the software. MD contributed to the software and analyzed the real data set. ET provided the probabilistic method to determine the minimizer threshold. KR designed the study and was a major contributor in writing the manuscript. All authors read and approved the final manuscript.

## Additional Files

Additional file 1 – Probabilistic Threshold Model

Contains additional information on how we obtained the minimizer thresholds used in the paper.

Additional file 2 – Command Line Arguments

Describes how we used the tools Mantis and COBS.

## References

1. Venter, J.C.,…, Reinert, K.,…, Zhu, X.: The sequence of the human genome. Science 291, 1304–1351 (2001)

2. International Human Genome Sequencing Consortium: Initial sequencing and analysis of the human genome. Nature 409(6822), 860–921 (2001)

3. Caulfield, M., Davies, J., Dennys, M., Elbahy, L., Fowler, T., Hill, S., Hubbard, T., Jostins, L., Maltby, N., Mahon-Pearson, J., McVean, G., Nevin-Ridley, K., Parker, M., Parry, V., Rendon, A., Riley, L., Turnbull, C., Woods, K.: The National Genomics Research and Healthcare Knowledgebase (2019). doi:10.6084/m9.figshare.4530893.v5

4. All of Us (NIH): All of Us Research Program Overview. (accessed: 15.09.2020) (2020). https://allofus.nih.gov/about/all-us-research-program-overview

5. Marchet, C., Boucher, C., Puglisi, S.J., Medvedev, P., Salson, M., Chikhi, R.: Data structures based on k-mers for querying large collections of sequencing datasets. bioRxiv, 866756 (2019). doi:10.1101/866756

6. Bingmann, T., Bradley, P., Gauger, F., Iqbal, Z.: COBS: A Compact Bit-Sliced Signature Index. In: String Processing and Information Retrieval, pp. 285–303. Springer, Cham (2019)

7. Berger, B., Peng, J., M.S.: Computational solutions for omics data. Nature reviews (2013)

8. Dadi, T.H., Siragusa, E., Piro, V.C., Andrusch, A., Seiler, E., Renard, B.Y., Reinert, K.: DREAM-Yara: an exact read mapper for very large databases with short update time. Bioinformatics (Oxford, England) 34(17), 766–772 (2018)

9. Wood, D.E., Salzberg, S.: Kraken: ultrafast metagenomic sequence classification using exact alignments. Genome biology 15(3), 46 (2014)

10. Solomon, B., Kingsford, C.: Fast search of thousands of short-read sequencing experiments. Nature Biotechnology 34(3), 300–302 (2016). doi:10.1038/nbt.3442. Accessed 2016-09-07

11. Sun, C., Harris, R.S., Chikhi, R., Medvedev, P.: AllSome Sequence Bloom Trees. bioRxiv, 090464 (2016). doi:10.1101/090464. Accessed 2017-03-08

12. Marchet, C., Iqbal, Z., Gautheret, D., Salson, M., Chikhi, R.: REINDEER: efficient indexing of k-mer presence and abundance in sequencing datasets. bioRxiv, 2020–0329014159 (2020). doi:10.1101/2020.03.29.014159

13. Pandey, P., Almodaresi, F., Bender, M.A., Ferdman, M., Johnson, R., Patro, R.: Mantis: A Fast, Small, and Exact Large-Scale Sequence-Search Index. Cell systems 7(2), 201–2074 (2018)

14. Bradley, P., den Bakker, H.C., Rocha, E.P.C., McVean, G., Iqbal, Z.: Ultrafast search of all deposited bacterial and viral genomic data. Nature Biotechnology 37(2), 152–159 (2019). doi:10.1038/s41587-018-0010-1

15. Gupta, G., Coleman, B., Medini, T., Mohan, V., Shrivastava, A.: RAMBO: Repeated And Merged Bloom Filter for Multiple Set Membership Testing (MSMT) in Sub-linear time, 1–14 (2019). 1910.02611

16. Holtgrewe, M.: Mason – a read simulator for second generation sequencing data. Technical Report FU Berlin (2010)

17. Kim, D., Song, L., Breitwieser, F.P., Salzberg, S.: Centrifuge: rapid and sensitive classification of metagenomic sequences. Genome research 26(12), 1721–1729 (2016)

18. Piro, V.C., Dadi, T.H., Seiler, E., Reinert, K., Renard, B.Y.: ganon: precise metagenomics classification against large and up-to-date sets of reference sequences. Bioinformatics (Oxford, England) 36(Supplement 1), 12–20 (2020)

19. Reinert, K., Dadi, T.H., Ehrhardt, M., Hauswedell, H., Mehringer, S., Rahn, R., Kim, J., Pockrandt, C., Winkler, J., Siragusa, E., Urgese, G., Weese, D.: The SeqAn C++ template library for efficient sequence analysis: A resource for programmers. Journal of Biotechnology (2017)

20. Marcais, G., Pellow, D., Bork, D., Orenstein, Y., Shamir, R., Kingsford, C.: Improving the performance of minimizers and winnowing schemes. Bioinformatics (Oxford, England) 33(14), 110–117 (2017)

21. Gog, S., Beller, T., Moffat, A., Petri, M.: From theory to practice: Plug and play with succinct data structures. In: 13th International Symposium on Experimental Algorithms, (SEA 2014), pp. 326–337 (2014)

22. Jokinen, P., Ukkonen, E.: Two algorithms for approximate string matching in static texts. Lecture Notes in Comp. Sci. 520(06), 240–248 (1991)

23. Bloom, B.H.: Space/time trade-offs in hash coding with allowable errors. Commun. ACM 13(7), 422–426 (1970). doi:10.1145/362686.362692

