## Additional File 1 for "Raptor: A fast and space-efficient pre-filter for querying very large collections of nucleotide sequences"

#### 0.1 Adapting the $k$-mer counting lemma for $(w, k)$-minimizers

In this section we address the extension and generalization of the $k$-mer lemma to (gapped) winnowing minimizers and present a probabilistic model to derive thresholds based on a user provided, tolerable loss rate and, as a single parameter, the number of minimizers. The thresholds can be precomputed and looked up during the filtering, improving the run time and filtering efficiency compared to ad hoc solutions. We provide the used data and a reference implementation on `https://github.com/eseiler/minimizer_thresholds`.

##### 0.1.1 Filtering with minimizers

For clarity we repeat some notations from the paper. We assume that we have a a query $p$ and a reference $T$ over an alphabet $\Sigma$. We will compare the query and the reference based on their sets of *representative* $k$-mers as described in the paper in Figure 1.

##### 0.1.2 A probabilistic model

In this section we model the distribution of minimizers that are affected by errors. In the following we refer to the size of the query as $p$. A shape $s$ is a subset of $\mathbb{N}$ representing the indices of the shape. For example, the shape #.##..# is represented by $s = \{0, 2, 3, 6\}$. The *size* of a shape is denoted by $k$ which is the cardinality of the set $s$. The *span* is denoted by $k' = \max s + 1$. Hence, for ungapped shapes the size equals the span, i.e. $k = k'$. The size of the window in which we compute minimizers is denoted by $w$ and $m$ is the number of minimizers in a query.

As in Lemma 1, an error affects several minimizers depending on its position and the character that is replaced. Each error lowers the threshold. We want to derive a probabilistic model that computes this threshold as good as possible with only $w, k, s$ and $m$ as input. We aim at having an as high as possible threshold without missing too many hits (i.e. having false negatives).

For this, we use the following definitions:

- $X_i$ : Random variable that is 1 if there is a minimizer starting at index $i$, $\forall i \in [0, p - k' + 1]$.
- $D_n$: Random variable that indicates that $n$ minimizers are affected by at least one error, $\forall n \in [0, p - k' + 1]$.
- $\tau$: Probability threshold for the number of affected minimizers.

**Index based model for one error**
In order to define $P(D_n)$ for one error, we need the distribution of $X_i$. In practice, assuming an uniform distribution of the minimizers in $[0,\ p - k' + 1]$ yields good results:

$$P(X_i) = \frac{\text{number of minimizer}}{\text{number of indices}} = \frac{m}{p - k' + 1} \tag{1}$$

This can now be used to compute $P(D_n)$. Let $j$ be the position within a sequence where the error is located. Then it can affect at most $k$ minimizers *directly*: The minimizers that are affected by the error at position $j$ are the minimizers starting at indices $j - s, \forall j \in s$. Naturally, these minimizers are the only ones that include the position $j$. Hence, there are at most $k$ minimizers that include the position $j$ and thereby there are at most $k$ minimizers *directly* modified by the error.
This can be expressed via a binomial distribution and yields:

$$\begin{aligned}
&\forall n \in [0, k] : \\
&P(D_n) = \binom{k}{n} P(X_i)^n (1 - P(X_i))^{k-n} \\
&\qquad = \binom{k}{n} \left(\frac{m}{p - k' + 1}\right)^n \left(1 - \frac{m}{p - k' + 1}\right)^{k-n}
\end{aligned} \tag{2}$$

$P(D_n)$ can now be used to compute the new filtering threshold. Let $d$ be the number of $\boldsymbol{(w, k, s)}$-minimizers affected such that

$$\sum_{i=0}^{d} P(D_i) > \tau \text{ and } \sum_{i=0}^{d-1} P(D_i) < \tau \tag{3}$$

This leads to the following lemma:

**Lemma 1** *For a given $\boldsymbol{p}$, $\boldsymbol{w}$, and $\boldsymbol{k}$ and one error, at most $\boldsymbol{d}$ many $\boldsymbol{k}$-mers are affected with a probability of $\boldsymbol{\tau}$. Thus, an approximate occurrence of the query in $\boldsymbol{T}$ has to share at least $\mathbf{t} = \boldsymbol{m} - \boldsymbol{d}$ $\boldsymbol{k}$-mers.*

**Lemma 1** can be applied to all of our 4 methods that are presented in this chapter. Notably, this method enables us to use $\boldsymbol{\tau}$ to control the false positive and false negative rates of our filter. The higher we choose $\boldsymbol{\tau}$, the more (gapped) $\boldsymbol{k}$-mers are affected, i.e. the threshold $\boldsymbol{t}$ decreases and the false positive rate increases. The lower we choose $\boldsymbol{\tau}$, the fewer (gapped) $\boldsymbol{k}$-mers are affected, i.e. the threshold increases $\boldsymbol{t}$ and the false negative rate decreases. Figures 4 and 5 show this effect with varying $\boldsymbol{\tau}$ for our final model.

Furthermore, this model only depends on $\boldsymbol{m}$ which is known for each read. In order to obtain the treshold, we simply have to look it up in a precomputed table for the specific parameters or compute the table once if it has not been computed yet.

##### Extension for indirect errors

The previous model can be extended to account for errors affecting minimizers *indirectly.* The example in Figure 3 shows that an error can affect a minimizer even though the error does not overlap with the minimizer.

We define the random variable $C_n$ that is 1 if the error affects $n$ $k$-mers indirectly. Thus, if we assume for simplification that the events of directly and indirectly affecting a minimizer are independent, $P(D_n)$ now becomes:

$$\begin{aligned}
&\forall n \in [0, w]: \\
&\qquad P(D_n) = \sum_{i=1}^{w-k} P'(D_{n-i}) P(C_i) \\
&\text{with} \\
&\qquad P'(D_n) = \begin{cases} P'(D_n) & \text{if } n <= k \\ 0 & \text{else} \end{cases}
\end{aligned} \tag{4}$$

where $P'(D_n)$ is the distribution of the previous model.

The main challenge with this extension is the computation of $P(C_n)$. We tried to find tractable formulation of $C_n$, but did not succeed and leave this as an open problem. $P(C_n)$ highly depends on the shape of the minimizers and the way they are computed, making $P(C_n)$ hard to precompute. This is why we estimate $P(C_n)$ in this work experimentally by sampling. We generate a query with and without an error and then check the number of minimizers that are indirectly modified by the error. This sampling method is flexible for different parameters and can be easily applied for the relevant range of parameters or be quickly computed on the fly. Note that $P(C_n)$ depends on the shape of the minimizers, the way they are computed, as well as the parameters $w$, $k$ and $m$. In our experiments, a sampling of 10,000 cases lead to convergence of $P(C_n)$.

**Extension for multiple errors**

We can further extend our model to consider multiple errors. For example, assume that there are 2 errors, $e_1$ and $e_2$, that affect $d_1$ and $d_2$ many $k$-mers, respectively. Thus, the two errors combined affect at most $d = d_1 + d_2$ many $k$-mers. Hence, affecting at most $d$ many $k$-mers can be achieved by all different combinations of $(d_1, d_2)$ such that $d \geq d_1 + d_2$ and $0 \leq d_1 \leq w$, $0 \leq d_2 \leq w$.

Generalizing this for $e$ errors affecting $d_1, \ldots, d_e$ minimizers leads to the following distribution:

$$\begin{aligned}
&\forall n \in [0, w \cdot e]: \\
&\qquad P(D_n) = \sum_{0 \leq d_1 \leq w, \ldots, 0 \leq d_e \leq w}^{d_1 + d_2 + \ldots + d_e = n} \prod_{i=1}^{e} P'(D_{d_i})
\end{aligned} \tag{5}$$

with $P'(D_n)$ being the distribution of the previous model.

##### Extension for overlapping errors

Figure 1 shows an example for overlapping errors. If two errors are close enough, they can affect the same $k$-mer which leads to an overestimation of affected minimizers by the current model.

```
a)(6,3,###)-minimizer  b)(6,3,###)-minimizer
    Without errors         With two errors
      TAAGCGGG               TAACTGGG
      -AAG--                  -AAC--
       AAG---                  AAC---
        AGC---                  ACT---
   2 ungapped 3-mers      2 ungapped 3-mers
```

Figure 1: **Example of two errors affecting the same $k$-mer.** a) and b) represent the same sequence without errors and with two errors at position 4 and 5 replacing a G with a C and a C with a T. Note that AAG has only one modification whereas AGC has two modifications but the previous method will count 3 modified $k$-mers when there are actually only 2 $k$-mers that have been modified.

We will refer to minimizers that are affected by at least $2$ errors as *overlapping* errors.

Incorporating overlapping errors into the previous model can be done by considering the probability of having $n$ errors that overlap, $\forall n \in [0, m]$. However, the exact computation of this probability requires to know all the different combinations of errors leading to $n$ overlapping errors. Furthermore, this cannot be simplified because the errors are not independent. In this section we will present a way to approximate the effect.

We assume that having more than $2$ errors *close* enough is unlikely to happen and thus only consider that $2$ errors lead to overlapping errors. Errors are close enough when the maximum distance between the positions of the errors is strictly lower than $k'$, hence the probability of these errors overlapping is greater than zero. For illustration we show the computation of this probability for the example in Figure 2.

```
(6,4,####)-minimizer
 -------------------
   ---- ----
    ----
     ----
      ----
       ----
```

Figure 2: **All the possible $k$-mers that are directly affected by two errors distinguished by overlapping.** — — —— represents a $k$-mer that is not affected by both errors and — — —— represents a $k$-mer that is affected by both errors. Note that for this specific error configuration there are 2 $k$-mers that are affected by both errors.

Figure 2 contains all the different $\boldsymbol{k}$-mers and highlights those two $\boldsymbol{k}$-mers that are affected by both errors. Thus, if these two $\boldsymbol{k}$-mers are minimizers, then there are two overlapping errors.
In our example, there are only 3 different cases: 0, 1 or 2 overlapping errors. The probability of each case can be easily computed by assuming that the minimizers are independent. With this assumption, the probability of having 0 overlapping error is the probability of having none of the *red* minimizers. Thus, the probability of this event is $\boldsymbol{(1-P(X_{e_1-1}))\cdot(1-P(X_{e_1}))}$ with $\boldsymbol{e_1}$ being the index of the first error. Using the same reasoning for 1 and 2 overlapping errors gives the probabilities $\boldsymbol{(1-P(X_{e_1-1}))\cdot P(X_{e_1})+P(X_{e_1-1})\cdot(1-P(X_{e_1}))}$ and $\boldsymbol{P(X_{e_1-1})\cdot P(X_{e_1})}$, respectively. The argumentation of this example can now be extended to all different configurations of $\boldsymbol{2}$ errors that are close enough.

Let $\boldsymbol{v(n)}$ be a function that approximates the probability of having $\boldsymbol{n}$ overlapping errors given the fact that there are $\boldsymbol{2}$ errors in a window. Hence, $\boldsymbol{v(n)}$ is simply the extension of the previous example to $\boldsymbol{n}$ overlapping errors without knowing the distance between 2 errors. Thus, $\boldsymbol{v(n)}$ can be computed by considering all the configurations where the distance between 2 errors can lead to $\boldsymbol{n}$ overlapping errors, i.e all the configurations where $\boldsymbol{e_2 - e_1 \leq k - n}$. After this, we need to consider the number of combinations that lead to $\boldsymbol{n}$ overlapping for each configuration. For a distance of $\boldsymbol{i = e_2 - e_1}$ there are exactly $\boldsymbol{\binom{k-n}{k-n-i} = \binom{k-n}{i}}$ possible overlapping errors. Each one of these overlapping errors occurs with a probability of $\boldsymbol{\bar{p}^n(1-\bar{p})^i}$. Thus, $\forall \boldsymbol{n} \in [\boldsymbol{0}, \boldsymbol{k-1}]$:

$$\boldsymbol{v(n) = \bar{p}^n \sum_{i=0}^{k-1-n} \binom{k-n}{i} (1-\bar{p})^i} \tag{6}$$

With $\boldsymbol{\bar{p} = P(X_n) = \frac{m}{p-k'+1}}$

To obtain a probability, we need to normalize:
$\forall\, [\boldsymbol{0}; \boldsymbol{k-1}]$,

$$\boldsymbol{P(O_n) = \frac{v(n)}{\sum_{i=0}^{k-1} v(i)}} \tag{7}$$

$\boldsymbol{P(O_n)}$ is the probability of having $\boldsymbol{n}$ overlapping errors.

This distribution can be used to remove the overlapping errors that are included in the $\boldsymbol{P(D_n)}$ previously computed:

$$\boldsymbol{P(D_n) = \sum_{o=0}^{min(k-1,w-n)} P(O_o)P'(D_{n+o})} \tag{8}$$

with $\forall \boldsymbol{n} \in [\boldsymbol{0}, \boldsymbol{w}]$, $\boldsymbol{P'(D_n)}$ the probability of the previous model.

Note that this extension is based on the assumption that errors are independent and that only $\boldsymbol{2}$ errors overlap. This is obviously not often the case, but for few errors the likelihood of it happening is very high. Moreover, note that this extension increases the threshold because it removes overlapping errors.

The previous model is based on $\boldsymbol{P(X_n)}$. However, this distribution has been assumed to be uniform thus leading to the fact that $\boldsymbol{P(X_n)} = \frac{\text{number of minimizer}}{\text{number of indices}}$. We tried to model this distribution more accurately, however all attempts lead to worse results than the uniform distribution.

##### 0.1.3 Using the probabilistic method

Our model includes one core method and two extensions that can be independently added. We will proceed as follows:

- Step 1: Compute $\boldsymbol{P(D_n)}$ for 1 error (more involved methods performed consistently worse).
- Step 2: Possibly extend the model for indirect errors (Choice I).
- Step 3: Compute $\boldsymbol{P(D_n)}$ for multiple errors.
- Step 4: Possibly extend the model for overlapping errors (Choice O).
- Step 5: Compute the threshold using $\boldsymbol{P(D_n)}$ as in Lemma 1.

Hence we have two independent choices:

- $\boldsymbol{I}$: The model takes indirect errors into consideration (Step 2).
- $\boldsymbol{O}$: The model takes overlapping errors into consideration (Step 4).

This means that there are 4 different ways to compute the threshold using this method. $\overline{\boldsymbol{X}}$ denotes not choosing $\boldsymbol{X}$, e.g., the model using ($\overline{\boldsymbol{I}}$,$\boldsymbol{O}$) skips Step 2 but performs Step 4.

In the following we aim to understand the differences between the 4 different methods and to compare them with ad hoc solutions.

In Figure 3 we computed the threshold of each method in function of the number of minimizers.

Figure 3 is giving us the expected results. Considering indirect errors lowers the threshold because we take more potentially affected minimizers into account. Overlapping errors do indeed increase the threshold since they reduce the number of affected minimizers by removing those that are counted twice. This figure also shows that the heuristic method is more strict and less consistent than our models. Note that the counting lemma threshold ($\boldsymbol{t = 150 - 26 + 1 - 3 \cdot 26 = 47}$) is not shown and that it is not directly comparable to our thresholds since we count minimizers while the counting lemma considers windows (which might contain the same minimizer).

To explore the true positive and true negative rates of our methods, we used the following data:

- A reference genome of size 10 million.
- $\boldsymbol{R_3}$: 100,000 reads of size $\boldsymbol{p \in \{100,\ 150,\ 250\}}$ which are extracted with 3 errors from the reference genome.

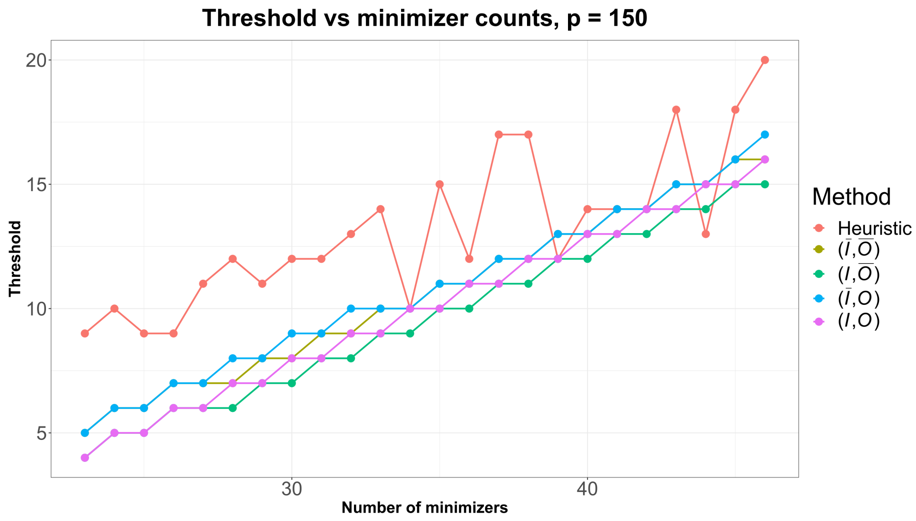


Figure 3: $p = 150$, $w = 26$, $k = 20$, $\tau = 0.99$. **Threshold of the different methods in function of the number of minimizers.** Colors denote the different methods. Note that the lemma threshold (47 independent of the number of minimizers) is not shown in the figure. While our models approximate a linear function, the heuristic method shows a huge variance.

- $R_{random}$: 100,000 reads that are randomly generated and hence should not match the reference.

Furthermore, we configured that:

- The number of errors $e$ is set to $3$.
- $k$-mers in the $(w, k, s)$-minimizers are ungapped, i.e $k = k'$.
- Minimizers in a window are selected using a lexicographically order.

We then computed for each read its $(w, k, s)$-minimizers and queried them in the reference data set's $(w, k, s)$-minimizers. If the number of shared $(w, k, s)$-minimizers (respectively the sum of the minimizers, weighted by their multiplicities, for the counting lemma) is higher than the threshold determined by the method, there is a match. We applied this procedure on the four different models, the counting lemma and the heuristic method.

Recall that our parameter $\tau$ is used to control the true positive and true negative rate. The true positive rate should be proportional to $\tau$.

Figure 4 highlights the fact that $\tau$ can be used to control the true positive rate. While a higher $\tau$ effects a higher true positive rate, it will also cause a lower true negative rate. Figure 5 shows that there are no false positives for $\tau$ greater than $0.95$ for our data.

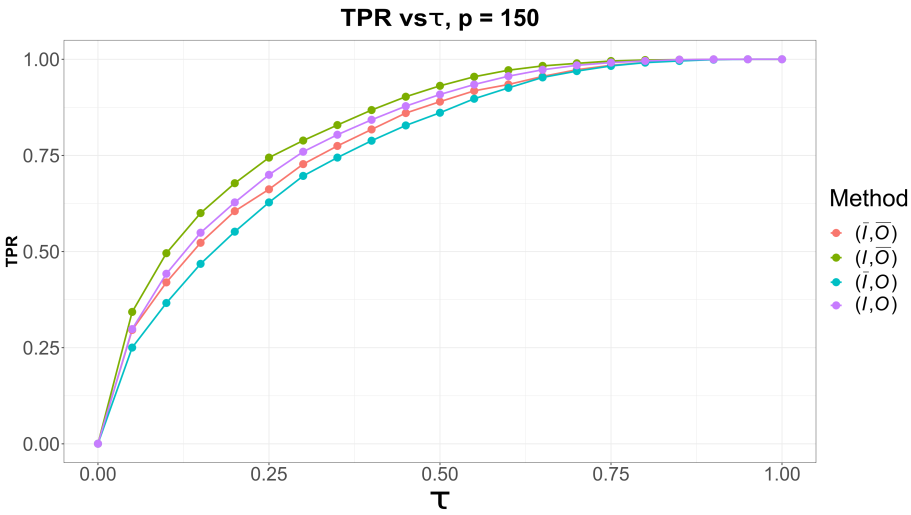


Figure 4: $\boldsymbol{p = 150}$**,** $\boldsymbol{w = 26}$**,** $\boldsymbol{k = 20}$**. True positive rate of the different models in function of** $\boldsymbol{\tau}$**.** Note that for $\boldsymbol{\tau > 0.85}$ we achieve a true positive rate of $\mathbf{1}$.

Note that for the values of $\boldsymbol{w}$, $\boldsymbol{k}$ and $\boldsymbol{p}$ of Figures 4 and 5, setting $\boldsymbol{\tau}$ to 0.95 will result in neither true negatives nor false positives.

To investigate the true positive and negative rates further, Figure 6 shows the F1-Score. As expected, there is wide range of $\boldsymbol{\tau}$ to choose from to achieve no false positives or negatives.

In Table 1 results for all methods can be seen. Notably, only the lemma and $\boldsymbol{(I, \overline{O})}$ find all reads. We can see that $\boldsymbol{I}$ decreases the threshold from 11.8 to 10.8 and adding $\boldsymbol{O}$ increases the threshold again to 11.3. The heuristic method yields an average threshold of 12.2 and hence finds fewer reads than the models. In Table 2 $\boldsymbol{\tau}$ was increased to 0.99. As a consequence, the average threshold of all models was reduced and the models found all reads.

The difference between our models and the heuristic and lemma method become clearer when approaching border cases, e.g., using reads of length 100 for our parameters as shown in Table 3. Here, the lemma returns a threshold of 0, hence all reads are reported as found, even the random reads. The heuristic method improves on that, however, still has $\mathbf{16,351}$ false positives and loses 3 true positives. Our models perform the best, reporting all true positive and reporting only 18 false positives. This shows that we perform well in the interesting border cases, where the thresholds are relatively low.

As a last measure, we want to compare the run times of the different methods. Table 4 shows the run times for querying reads of length 100 in the reference. Our models are as fast as the lemma when comparing the time to compute the threshold and are about 200 times faster than the heuristic method in that regard. Both Lemma and heuristic method do not need to conduct precompu-

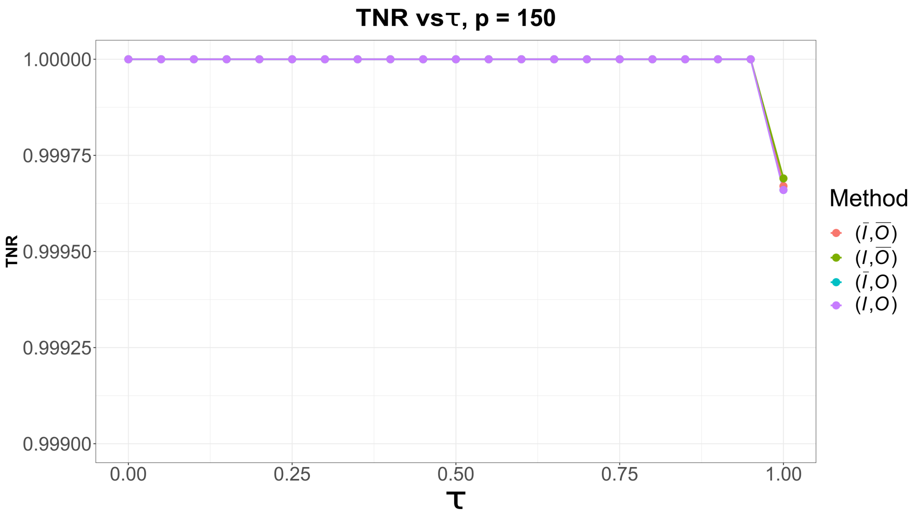


Figure 5: $p = 150$**,** $w = 26$**,** $k = 20$**. True negative rate of the different models in function of** $\tau$**.** Note the different scale of the y-axis. For a $\tau$ up to $0.95$ we generate no false positives.

tations - a necessary step for our models that takes around 1000 times (approx. 4 seconds) longer than the actual threshold computation. If the computations have been done once, the results are stored to disk and can be reused. In this case, the precomputation only consists of reading the file and hence the time for precomputation becomes negligible.

Moreover, note that the value of $\tau$ does not impact the computation time because increasing $\tau$ only results in slightly increasing the number of additions required to compute $d$ as defined in the previous section.

In conclusion, our approach based on a probabilistic model is able to accurately represent the number of minimizers affected by multiples errors. To do that, the user can select the appropriate extensions $I$ and $O$ to fit his need. Previous results yield the conclusion that, depending on the parameters, results of the models can slightly vary.

We suggest the following way to determine the best model: The value of $p$, $w$, $k$, $e$ already have been chosen. So all that is left is $\tau$. The next step is to estimate the lowest number of minimizers expected for the selected parameters. If this number is *low*, then consider using $(\overline{I},\overline{O})$ or $(\overline{I},O)$, because they lead to higher thresholds. If the threshold is *very low* then consider using a $\tau \leq 0.99$. In all other cases, $(I,O)$ might be the best option. $\tau = 0.99$ should be enough to guarantee few false negatives and positives.

Our models are especially powerful when we are expecting many $k$-mers to be destroyed by errors. For example, the lemma collapses, i.e. returns 0, when searching for 3 errors in a pattern of length 100 and a windows size of 26, thus providing no filtering. While the heuristic method performs better than the

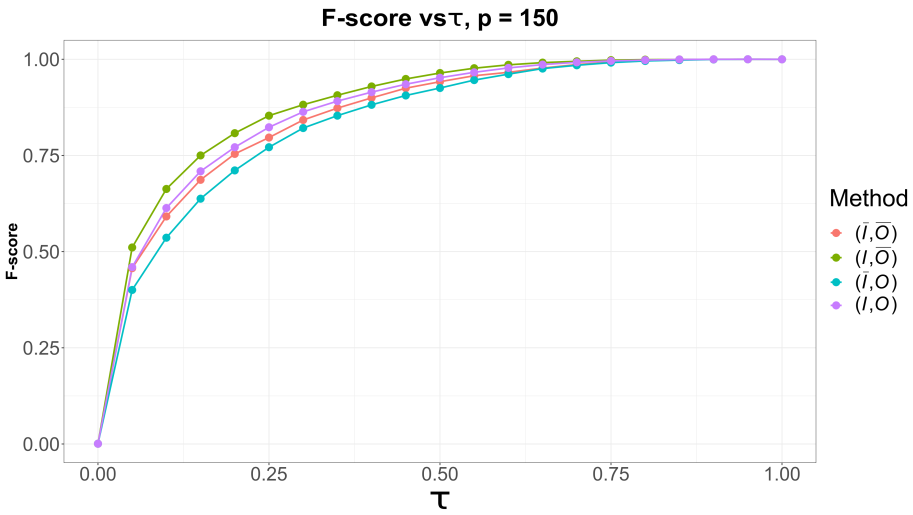


Figure 6: $\boldsymbol{p = 150}$, $\boldsymbol{w = 26}$, $\boldsymbol{k = 20}$. **F1-Score of the different models in function of** $\boldsymbol{\tau}$**.** Note that there is a region around $\boldsymbol{\tau = 0.95}$ where the F1-Score is $\mathbf{1}$.

lemma, it performs worse than our models.

Note that the value of $\boldsymbol{\tau}$ needs to be adjusted to your needs. The higher $\boldsymbol{\tau}$ is, the fewer false negatives are to be expected.

Moreover, the strength of this approach is that it does not depend on the reads itself. It only relies on known information such as $\boldsymbol{p}$, $\boldsymbol{w}$, $\boldsymbol{k}$, $\boldsymbol{s}$, the way minimizers are chosen, etc. This allows the thresholds to be precomputed, making this method easy to use and also really fast. Our models' threshold only depends on $\boldsymbol{m}$, the number of minimizers of the reads. That means that some thresholds are computed but never used. A simple way to avoid this problem is to not precompute the thresholds but to compute them on the fly and to store the threshold each time a read has a new number of minimizers $\boldsymbol{m}$.

### References

| Method | $\boldsymbol{R_3}$ | $\boldsymbol{R_{random}}$ | min threshold | average threshold |
|---|---|---|---|---|
| ($\overline{\boldsymbol{I}}$,$\overline{\boldsymbol{O}}$) | 99,996 | 0 | 7 | 11.8 |
| ($\boldsymbol{I}$,$\overline{\boldsymbol{O}}$) | 100,000 | 0 | 7 | 10.8 |
| ($\overline{\boldsymbol{I}}$,$\boldsymbol{O}$) | 99,995 | 0 | 8 | 11.8 |
| ($\boldsymbol{I}$,$\boldsymbol{O}$) | 99,998 | 0 | 7 | 11.3 |
| Heuristic | 99,991 | 0 | 3 | 12.2 |
| Lemma | 100,000 | 0 | 47 | 47.0 |

Table 1: **$\boldsymbol{p = 150}$, $\boldsymbol{w = 26}$, $\boldsymbol{k = 20}$, $\boldsymbol{\tau = 0.95}$. Number of reads selected in function of the data set.** $\boldsymbol{R_3}$ represents true positive queries and $\boldsymbol{R_{random}}$ represents false positive queries. We also show the minimum and average threshold returned by the respective method. Note that the threshold of the lemma is not directory comparable to the others since it takes multiplicity in account and considers windows instead of minimizers.

| Method | $\boldsymbol{R_3}$ | $\boldsymbol{R_{random}}$ | min threshold | average threshold |
|---|---|---|---|---|
| ($\overline{\boldsymbol{I}}$,$\overline{\boldsymbol{O}}$) | 100,000 | 0 | 5 | 9.3 |
| ($\boldsymbol{I}$,$\overline{\boldsymbol{O}}$) | 100,000 | 0 | 4 | 8.3 |
| ($\overline{\boldsymbol{I}}$,$\boldsymbol{O}$) | 100,000 | 0 | 5 | 9.6 |
| ($\boldsymbol{I}$,$\boldsymbol{O}$) | 100,000 | 0 | 5 | 9.6 |
| Heuristic | 99,991 | 0 | 3 | 12.2 |
| Lemma | 100,000 | 0 | 47 | 47.0 |

Table 2: **$\boldsymbol{p = 150}$, $\boldsymbol{w = 26}$, $\boldsymbol{k = 20}$, $\boldsymbol{\tau = 0.99}$. Number of reads selected.** $\boldsymbol{R_3}$ represents true positive queries and $\boldsymbol{R_{random}}$ represents false positive queries. We also show the minimum and average threshold returned by the respective method. Note that the threshold of the lemma is not directory comparable to the others since it takes multiplicity in account and considers windows instead of minimizers.

| Method | $\boldsymbol{R_3}$ | $\boldsymbol{R_{random}}$ | min threshold | average threshold |
|---|---|---|---|---|
| ($\overline{\boldsymbol{I}}$,$\overline{\boldsymbol{O}}$) | 100,000 | 18 | 1 | 1.0 |
| ($\boldsymbol{I}$,$\overline{\boldsymbol{O}}$) | 100,000 | 18 | 1 | 1.0 |
| ($\overline{\boldsymbol{I}}$,$\boldsymbol{O}$) | 100,000 | 18 | 1 | 1.0 |
| ($\boldsymbol{I}$,$\boldsymbol{O}$) | 100,000 | 18 | 1 | 1.0 |
| Heuristic | 99,997 | 16,351 | 0 | 1.6 |
| Lemma | 100,000 | 100,000 | 0 | 0.0 |

Table 3: **$\boldsymbol{p = 100}$, $\boldsymbol{w = 26}$, $\boldsymbol{k = 20}$, $\boldsymbol{\tau = 0.99}$. Number of reads selected.** $\boldsymbol{R_3}$ represents true positive queries and $\boldsymbol{R_{random}}$ represents false positive queries. We also show the minimum and average threshold returned by the respective method. Note that our models produce way fewer false positives than the heuristic or lemma approach.

| Method | Precomputation time | | Thresholding time |
|---|---|---|---|
| | Compute | Load | |
| $(\bar{\boldsymbol{I}},\bar{\boldsymbol{O}})$ | 4177 | 0.08 | 4.4 |
| $(\boldsymbol{I},\bar{\boldsymbol{O}})$ | 4258 | 0.1 | 4.2 |
| $(\bar{\boldsymbol{I}},\boldsymbol{O})$ | 4110 | 0.09 | 4.2 |
| $(\boldsymbol{I},\boldsymbol{O})$ | 4222 | 0.09 | 5 |
| Heuristic | | | 868.2 |
| Lemma | | | 4.5 |

Table 4: **$p = 100$, $w = 26$, $k = 20$, $\tau = 0.99$. Run times of the different methods in milliseconds.** The precomputation time only applies to the computation of the models. This can be done by either computing on the fly or loading the results from a file. Thresholding encompasses the selection of minimizers and the method-specific operation, i.e. look-up of the threshold, applying the heristic method, or computing the lemma.
