## Additional File 2 for "Raptor: A fast and space-efficient pre-filter for querying very large collections of nucleotide sequences"

#### 0.1 Used Command Line Arguments for Comparison with Mantis and COBS

For the artificial data set, COBS was built with the arguments $compact - construct$, $k = 19$, --*num-hashes* 2 and -*t* 32. All other arguments, especially the FPR, were used with their default values.
For the real data set, COBS was built with the arguments $compact - construct$ and $k = 20$, all other arguments were used with their default values.
Mantis was built on top of Squeakr files. The Squeakr files were constructed by using the flag -*e* for exact construction, the flag -*n* to store only the necessary information. -*k* was set to 19 for the artificial data sets and to 20 for the real data set. Moreover, for the real data set a cutoff based on the file size of the gzipped FASTQ file was set by using the argument -*c*. The cutoffs Mantis used were proposed by Solomon and Kingsford [2] and were also used for the IBF construction, the exact cutoffs can be found in 0.1.

| **Min Size** | **Max Size** | **Cutoff** |
|---|---|---|
| 0 | $\leq$ 300MiB | 1 |
| ¿ 300MiB | $\leq$ 500MiB | 3 |
| ¿ 500MiB | $\leq$ 1GiB | 10 |
| ¿ 1GiB | $\leq$ 3GiB | 20 |
| ¿ 3GiB | $\infty$ | 50 |

Table 1: Cutoffs

Mantis was then run with the parameter -*s* set to 34, as recommended by [1]. After the build, the mantis index was further compacted by running *mantis mst*. The reported construction time for Mantis was determined by adding the running time of both processes.
For the construction of the IBFs for the real data set, the build function was first run using the flag --*compute-minimiser*, which precomputes minimizer files. Then the actual IBF is constructed by running the build function with the precomputed minimizers. The reported construction time is the run time of this second build call.
Furthermore, like COBS on the real data set, only one hash function was used for the IBFs. The size has to be set beforehand and was determined to be 8 GiB for the $(20, 20)$-IBF and 800 MiB for the $(40, 20)$-IBF.

### References


[1] P. Pandey, F. Almodaresi, M. A. Bender, M. Ferdman, R. Johnson, and R. Patro. Mantis: A Fast, Small, and Exact Large-Scale Sequence-Search Index. *Cell systems*, 7(2):201–207.e4, Aug. 2018.

[2] B. Solomon and C. Kingsford. Fast search of thousands of short-read sequencing experiments. *Nature Biotechnology*, 34(3):300–302, Mar. 2016.
